# A modular dCas9-based recruitment platform for combinatorial epigenome editing

**DOI:** 10.1101/2022.07.01.498378

**Authors:** Tessa Swain, Christian Pflueger, Saskia Freytag, Daniel Poppe, Jahnvi Pflueger, Trung Nguyen, Ji Kevin Li, Ryan Lister

**Author notes:** These authors contributed equally (Tessa Swain, Christian Pflueger). The authors wish it to be known that, in their opinion, the first two authors should be regarded as joint First Authors.

## Abstract

CRISPR-dCas9 based targeted epigenome editing tools allow precise manipulation and functional investigation of various genome modifications. However, these tools often display substantial context dependency, with highly variable efficacy between target genes and cell types, potentially due to underlying variation in the chromatin modifications present. While simultaneous recruitment of multiple distinct ‘effector’ chromatin regulators has improved efficacy, these systems typically lack control over which effectors bind and their spatial organisation. To overcome this we have created a new modular combinatorial epigenome editing platform, called SSSavi. This system acts as an interchangeable and reconfigurable docking platform fused to dCas9 to enable simultaneous recruitment of up to four different effectors, allowing precise control and reconfiguration of the effector composition and spatial ordering of their binding. We demonstrate the activity and specificity of the SSSavi system and compare it to existing multi-effector targeting systems, establishing its efficacy. Furthermore, by altering the spatial ordering of effector recruitment, across multiple target genes and cell lines, we demonstrate the importance of effector recruitment order for effective transcriptional regulation. Together, this system offers the capacity to explore effector co-recruitment to specific loci to potentially enhance the manipulation of chromatin contexts previously resistant to targeted epigenomic editing.

## INTRODUCTION

The epigenome consists of chemical modifications to DNA and histone proteins that act as supplementary layers of information on the genome that can alter transcriptional activity, chromatin organisation, and DNA accessibility [1,2]. In recent years, genome-wide analyses have provided many new insights into the genomic distributions, dynamics, cell-type specificity, and combinatorial patterns of a range of epigenome modifications, allowing associative inferences of the role of these marks in transcriptional regulation [3,4]. However, it has been challenging to establish the precise roles of these multifarious modifications due to the natural complexity of the many different combinations and genomic contexts they are found in [5]. The gap in knowledge between the correlative data and the causal functions of epigenome modifications is a major limitation in both understanding and manipulating these important regulatory layers of the genome. Therefore, the ability to effectively edit the epigenome to investigate and understand its causal effects upon transcription, and to correct improper epigenetic states, will open up useful applications for biotechnology and synthetic biology. Early attempts to drive a targeted change in the epigenome fused programmable DNA binding domains to epigenome modifying ‘effector’ proteins to a desired target locus, achieving changes in a variety of modifications and gene expression [6–10]. However, these tools suffer from major limitations: generally, they target only a single epigenome modification and in mosts cases always fail to induce stable changes [11], they are frequently ineffective in altering gene expression or chromatin state [11,12], and they show highly variable efficacy depending on target genes and cell types [13]. A major challenge that likely underpins the limited performance of these tools is the naturally occurring combinatorial complexity of epigenome modifications and regulatory pathways [14]. Currently, only a single effector or very limited combinations have been tested for their role in epigenome editing and transcriptional activation [7,15–17]. Additionally, although some effective combinations of transcriptional repressors have been identified [6,18], they also fail to perform in all contexts, and utilise a direct fusion design that limits flexibility in terms of altering the effectors recruited. Thus, further improvement of these tools would be valuable to address the issue of limited efficacy as well as genomic context and cell type specific effects.

Recent studies assessing combinations of chromatin modifying effectors have examined their effects by fusing them directly to a DNA binding protein such as dCas9. Several studies [6,9,11,18–21] have reported enhanced, stable transcriptional regulation that was dependent upon the co-recruitment of combinations of effectors directly fused to the DNA binding protein [6,9,11,18–21]. These direct fusion approaches have revealed novel regulatory behaviours distinct from using components in isolation. A critical consideration is that some factors may have obligate binding partners or require co-localisation of other regulator enzymes, and thus when assessed individually may appear non-functional or display limited efficacy [22]. Thus, the above work highlights the need to expand testing of more complex interactions to determine their combinatorial regulatory properties. Importantly, there are hundreds of predicted epigenome regulatory proteins encoded in the human genome [23]. Thus, this provides an opportunity to further expand and advance the repertoire of epigenome editing tools to more effectively manipulate the readout of genomic information, and to understand the hierarchies and interactions between different modifications. However, the direct fusion constructs in the aforementioned studies limit the flexibility to include new effectors and test more than simple interactions.

Moving beyond the early systems that fuse a single effector to dCas9 [24–28], a range of more versatile platforms have been developed that allow recruitment of multiple effectors to dCas9 and the target locus, including the RNA aptamer assembly-based SAM platform [29], the SunTag system [30], or coiled-coil heterodimer pairs [31]. However, these systems have several limitations. RNA aptamers such as MS2 and PP7 are restricted by both the number of aptamers available and the number that can be included in a sgRNA [29,32,33], thereby constraining effector recruitment. Protein tags fused to dCas9 can offer greater flexibility. The SunTag system, for example, can harbour up to 24 GCN4 peptide repeat docking sites, thus providing an epitope binding site for an equivalent number of effectors [30]. However, the SunTag is limited to the same single epitope-antibody interaction for each docking site on the GCN4 peptide array. Consequently, when recruiting multiple distinct effector proteins fused to the *α*GCN4 domain, each will compete for binding to the SunTag, thereby limiting stoichiometric control and not allowing effector binding order to be precisely programmed. Such control can be important for the efficacy of combinatorial systems, as first demonstrated for the multicomponent VPR system [7]. Thus to overcome these limitations, here we report the creation of a new multi-site docking platform for epigenome editing, which we term the Spy-Snoop-Sun-Avi (“SSSavi”) system. This new platform has the highly valuable ability to control the order in which different effectors are recruited to a target site, while also allowing effector composition and stoichiometry to be easily changed, thus providing a means of addressing multiple current shortcomings of combinatorial epigenome editing systems.

## RESULTS

### Selection and justification of the four SSSavi tags

To overcome the limitations in the compositional versatility of existing epigenome editing tools, we aimed to construct and test a new epigenome editing platform that contains four distinct protein-protein interaction docking domains. This “SSSavi” system, fused to dCas9, is designed to recruit interchangeable combinations of up to four different effectors in a highly specific manner while allowing precise control over the ordering in which they bind. This was achieved by utilising four distinct protein-protein interaction domains, the Spy [34], Snoop [35], Sun [30] and Avi tags [36] (hence SSSavi) and their corresponding binding partners (herein referred to as catchers, **Figure 1A**). The Spy and Snoop tags were selected as they each spontaneously recombine with their counterpart catcher domains to form covalent bonds within minutes, while demonstrating no cross-reactivity between the Spy and Snoop components [34,35]. The SunTag [30] has been widely adopted as a modular recruitment platform [37–42] and consists of up to 24 short GCN4 peptides, each of which acts as epitope docking site for its counterpart mono-chain antibody, scFvGCN4 (*α*GCN4). By fusing the SunTag array to a programmable DNA binding protein such as dCas9, while fusing the counterpart antibody *α*GCN4 to an effector, one can attain amplified recruitment of the effector to desired DNA target sites in the genome. The fourth component used for the SSSavi docking array was a streptavidin variant known as Traptavidin [36] was selected as it has a superior biotin-binding stability compared to streptavidin [36]. Introduction of the *E. coli* biotin ligase, BirA, allows for targeted covalent biotinylation of the short AviTag [43]. Thus, these four highly stable binding pairs provide four unique docking sites that are the foundation of the SSSavi system.

**Figure 1.**
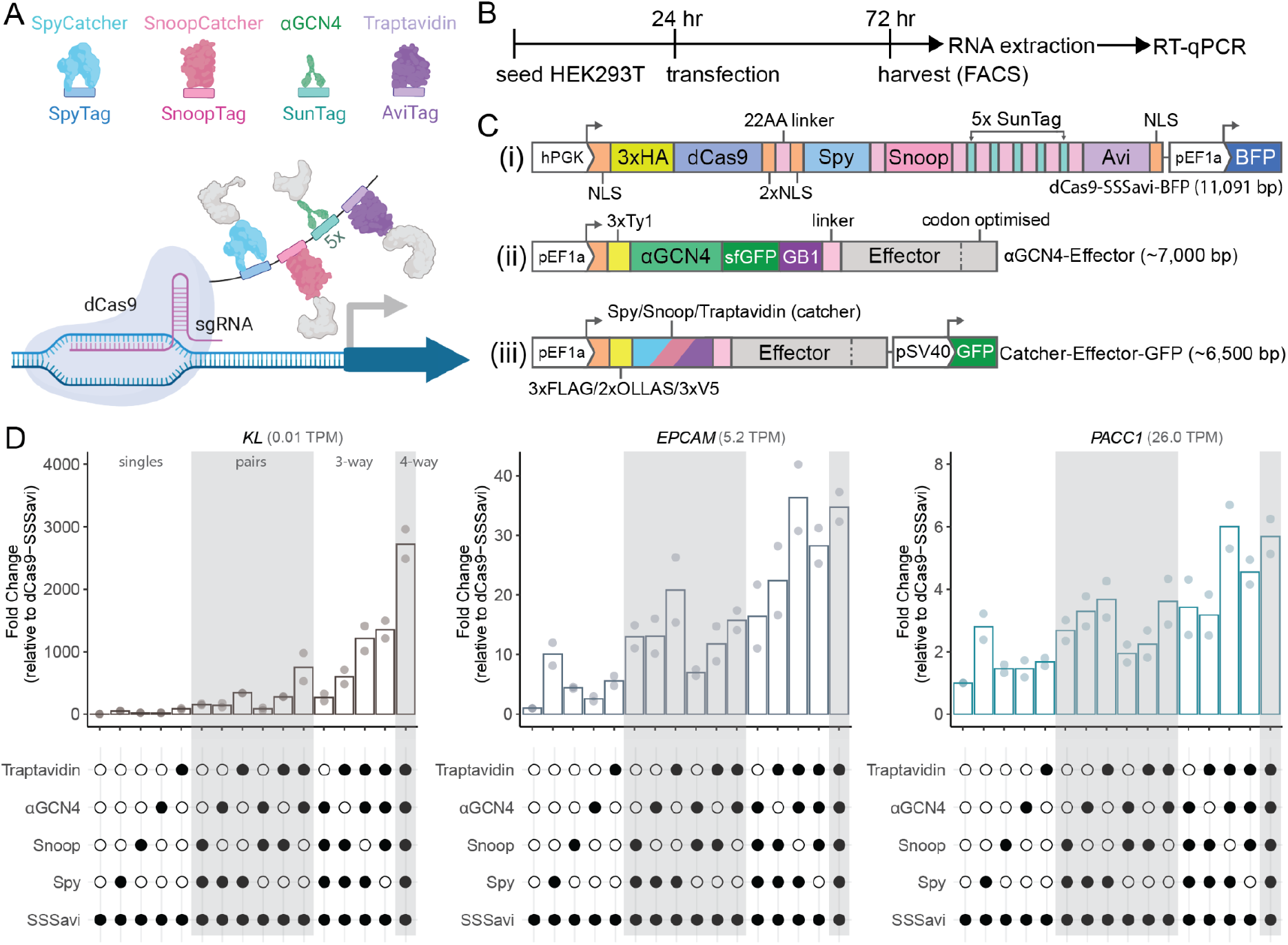
Successive addition of effectors using the SSSavi docking platform results in amplified transcriptional activation. **(A)** A schematic of the SSSavi docking system consisting of four tags (Spy, Snoop, Sun, and Avi) and their counterpart binding partners, and interaction of Spy catcher, Snoop catcher, *α*GCN4, and Traptavidin with a locus via fusion of the SSSavi docking array to dCas9. **(B)** Outline of experimental setup and timeline for construct delivery and measuring changes in gene expression. **(C)** Schematic of SSSavi plasmid constructs including **(i)** dCas9-Spy-Snoop-Sunx5-AviTag-BFP (dCas9-SSSavi-BFP), **(ii)** *α*GCN4 with internal sfGFP and GB1 (solubilising factor) domains, and **(iii)** the Spy, Snoop, and Traptavidin catcher-effector constructs with a separate GFP reporter. Constructs not to scale. **(D)** RT-qPCR quantification of *KL, EPCAM* and *PACC1* target gene expression following construct transfection, comparing recruitment to the SSSavi docking array and target gene promoters of one-, two-, three-, and four-way combinations of the different catcher domains linked to the super activator p65HSF1 (n = 2, biological replicates, bar graphs indicate sample mean, reference sample: dCas9-SSSavi only transfected cells). Cells tested were HEK293T 6x sgRNA stable cells (H6G), sorted for BFP (dCas9) and GFP (catchers).

### Design and testing of the SSSavi platform

Testing of multiple components and design features was required to determine the most optimal configuration for effective recruitment of effectors and induction of changes in gene expression. Design considerations included: (1) whether the smaller tag or larger catcher domain should be fused to the effector; (2) which linker sequences should be used for the SSSavi docking array and effector fusions; and (3) whether single or multiple binding domains should be used in the docking array.

To determine whether the docking array should be constructed from the smaller tags or the larger catcher domains we tested which was more effective for targeted transcriptional activation of a panel of endogenous genes in HEK293T cells. The super activator p65HSF1 [29] was fused to either the tag or the catcher, and co-transfected with the counterpart tag or catcher fused to dCas9 (**Supplementary Figure S1A**). These constructs were transfected into the H6G cell line, which is a HEK293T cell line that stably expressess six distinct sgRNAs targeting the promoters of *KL, EPCAM, PACC1, B2M, RBM3*, and *HINT1* (**Supplementary Figure S2A**, the best performing sgRNA (red) was selected for stable integration based on the screening of up to 6 sgRNAs per target promoter, **Supplementary Figure S2B**). These 6 target genes exhibit a wide range of transcript levels (0 - 716 Transcripts Per Million, TPM), allowing the regulatory capacity of different effectors to be tested at promoters of different strengths. Transfected cells were collected 72 h post-transfection (ptf) via fluorescence-activated cell sorting (FACS), followed by RT-qPCR (**Figure 1B**). Target gene activation was similar when either the tag or the catcher was fused to the p65HSF1 transcriptional activator (**Supplementary Figure S1B**). The smaller dCas9-SpyTag and dCas9-SnoopTag fusions marginally outperformed the larger dCas9-catcher fusion constructs when targeted to *EPCAM* (*p* = 0.01 and *p* = 0.03, respectively). Consequently, further testing used the dCas9-tag fusions, and use of the tags to construct the docking array.

The second design feature to be considered was the linker sequences. Starting with the SSSavi docking array, each of the four tags (Spy, Snoop, Sun and Avi) was separated by a 22 amino acid linker sequence, as previously used in one SunTag implementation [37] (**Figure 1C**). This longer linker was selected with the aim of more effectively recruiting multiple different effectors of varying sizes, compared to the shorter five amino acid linkers used by Tanenbaum *et al*. [30]. For the linkers used to fuse catcher domains to effectors, the Spy, Snoop, and Avi catchers linkers mimick that of the original *α*GCN4 fusion in the SunTag (**Figure 1C**), while excluding the sfGFP and GB1 domains (**Figure 1C**), as these potentially reduced protein stability (**Supplementary Figure S3**). A pSV40-GFP reporter was added to the catcher plasmids to allow FACS based enrichment of transfected cells, while the 3’ 300 bp of each effector was codon optimised in order to distinguish their expression from any endogenous counterpart effectors.

Finally, while a single domain of the Spy, Snoop, and Avi tags were used in the SSSavi platform, we incorporated a 5xSunTag array, rather than only 1xSunTag domain, as little to no target gene activation was observed from the latter when recruiting *α*GCN4-p65HSF1, compared to the use of 5xSunTag repeats (**Supplementary Figure S4**). Thus answering the question for the third design feature, whether one or more tags should be utilised to achieve effective effector recruitment.

### Combinatorial recruitment and specificity of the SSSavi system

Initial testing was undertaken to establish the regulatory capacity of the complete SSSavi docking array by fusing p65HSF1 to the C-terminus of each catcher and subsequent transfection into H6G cells. The ability of each single catcher to upregulate the three most lowly expressed target genes was examined using RT-qPCR, along with testing each set of two-, three-, and four-way catcher-p65HSF1 combinations (**Figure 1D**). Overall, the successive addition of the different catcher-p65HSF1 components resulted in higher transcriptional activation when recruiting multiple copies of p65HSF1 to the SSSavi docking array and target promoter. This effect was similar across all three target genes examined, despite the large variation in their endogenous expression levels. Specifically, the SSSavi system recruiting all four catcher-p65HSF1 domains resulted in a 2722-fold, 35-fold and 5.7-fold increase for *KL, EPCAM*, and *PACC1*, respectively (*p*-values are provided in **Supplementary Table S3**).

We next tested for possible cross-reactivity, or non-specific interactions, between the dCas9-tag:catcher components to determine whether the transcriptional upregulation was a result of specific tag:catcher binding events. Each dCas9-tag and catcher was tested for possible cross-reactivity by transfecting H6G cells with one of the following dCas9 constructs: dCas9-Spy, dCas9-Snoop, dCas9-SunTag, or dCas9-Avi, in conjunction with one of four catchers fused to p65HSF1 (**Figure 2A**). Quantitation of transcript abundance at multiple target genes (**Figure 2B**) demonstrated that all cognate tag:catcher pairs caused significant upregulation (*KL*: 452-881x, *EPCAM*: 7-15x, *PACC1*: 2-3x) compared to when there was no catcher (SpyTag or AviTag only), and all cognate pairs caused activation compared to non-cognate pairs. While a subset of the non-cognate pairs cause some limited upregulation compared to when there was no catcher (*KL*: 13-68x, *EPCAM*: 1.6-3.1x, *PACC1*: 1.2-1.4), this was significantly lower than that achieved by the cognate pairs, demonstrating the specificity of the tag:catcher recruitment.

**Figure 2.**
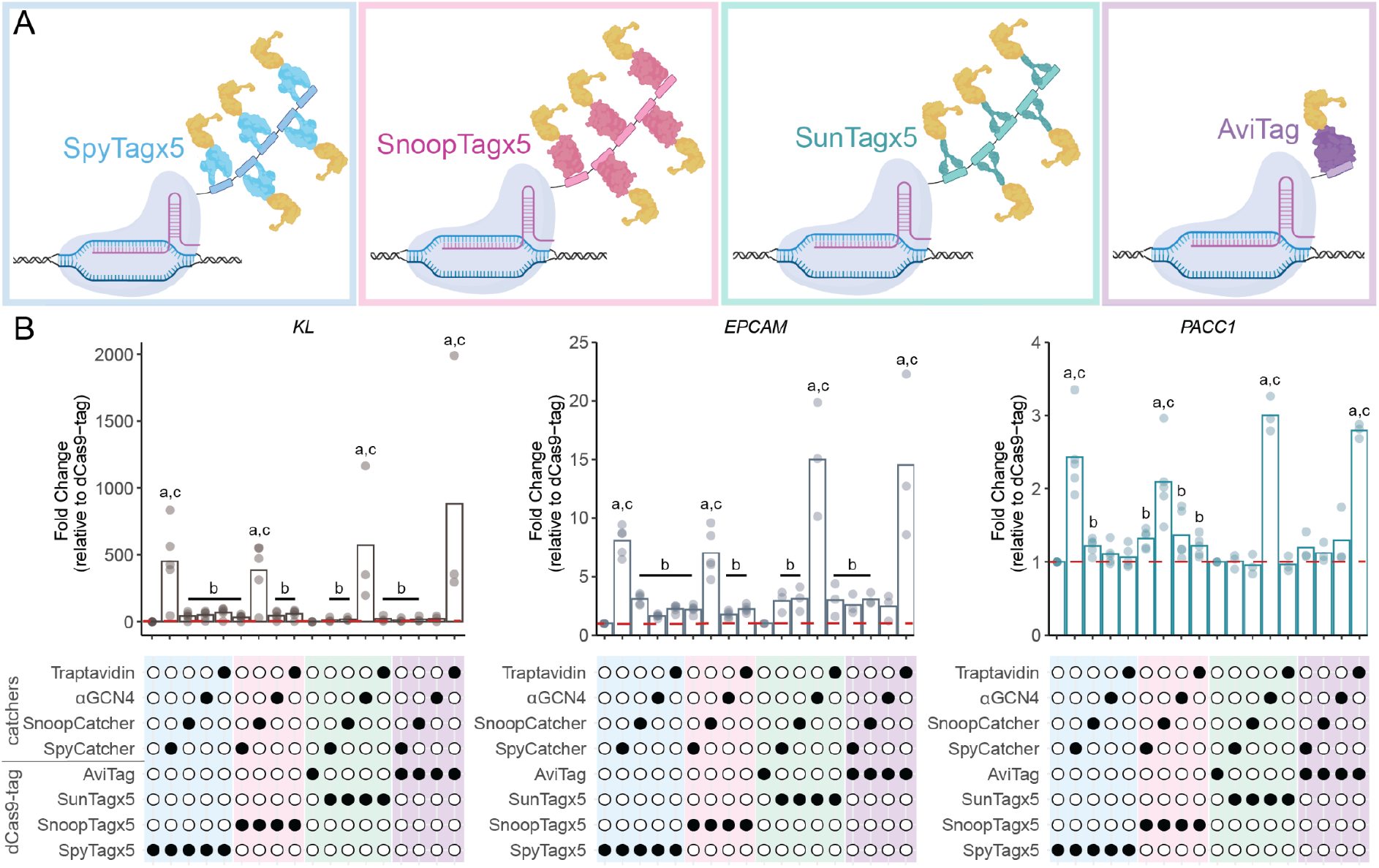
Binding specificity of tags and catchers. **(A)** A schematic of the four dCas9-tag platforms tested: SpyTagx5 (blue), SnoopTagx5 (pink), SunTagx5 (green) and AviTag (purple), each recruiting p65HSF1. **(B)** RT-qPCR quantification of target gene transcriptional activation for different dCas9-tag:catcher combinations. Each dCas9-tag construct was co-transfected with either SpyCatcher, SnoopCatcher, *α*GCN4, or Traptavidin fused to p65HSF1 (n = 3-5, biological replicates, bar graphs indicate sample mean, reference condition: dCas9-Tag (Spy or Avi as indicated) with baseline fold change value of 1 indicated by red dotted line). Cells tested were HEK293T 6x sgRNA stable cells (H6G), sorted for BFP and GFP. Bars with different letters indicate a significant difference as calculated by independent sample t-tests with Benjamini-Hochberg correction when comparing (a) each cognate tag:catcher pair to no catcher (SpyTag or AviTag only), (b) each non-cognate pair to no catcher, and (c) each cognate pair to non-cognate pair (*p*-values < 0.05).

### Comparison of SSSavi to commonly used dCas9-based activation systems

Having established the specificity and efficacy of the SSSavi system components, we next assessed the level of transcriptional activation achieved with the SSSavi docking array compared to well-established and commonly used activation systems including dCas9-activator direct fusions (-p65HSF1, VPR, VP64), and dCas9-SunTagx5. This was done by transiently transfecting each of the different activation systems into H6G cells, along with a transcriptional activator, as illustrated in **Figure 3A**. This demonstrated that the SSSavi system activates comparably, if not better, than other dCas9 recruitment platforms, with a notable gene-specific effect visible for dCas9-VPR when recruited to *PACC1* (**Figure 3B**). A single Spy or Snoop catcher fused to p65HSF1 performed comparably to five copies of VP64 recruited to the SunTag when targeting *EPCAM* and *PACC1*, while outperforming the dCas9-VP64 and -p65HSF1 direct fusions for *EPCAM*. This effect is more pronounced for the *α*GCN4 and Traptavidin components, as they are comparable if not better performers than even the VPR system for *KL* and *EPCAM*.

**Figure 3.**
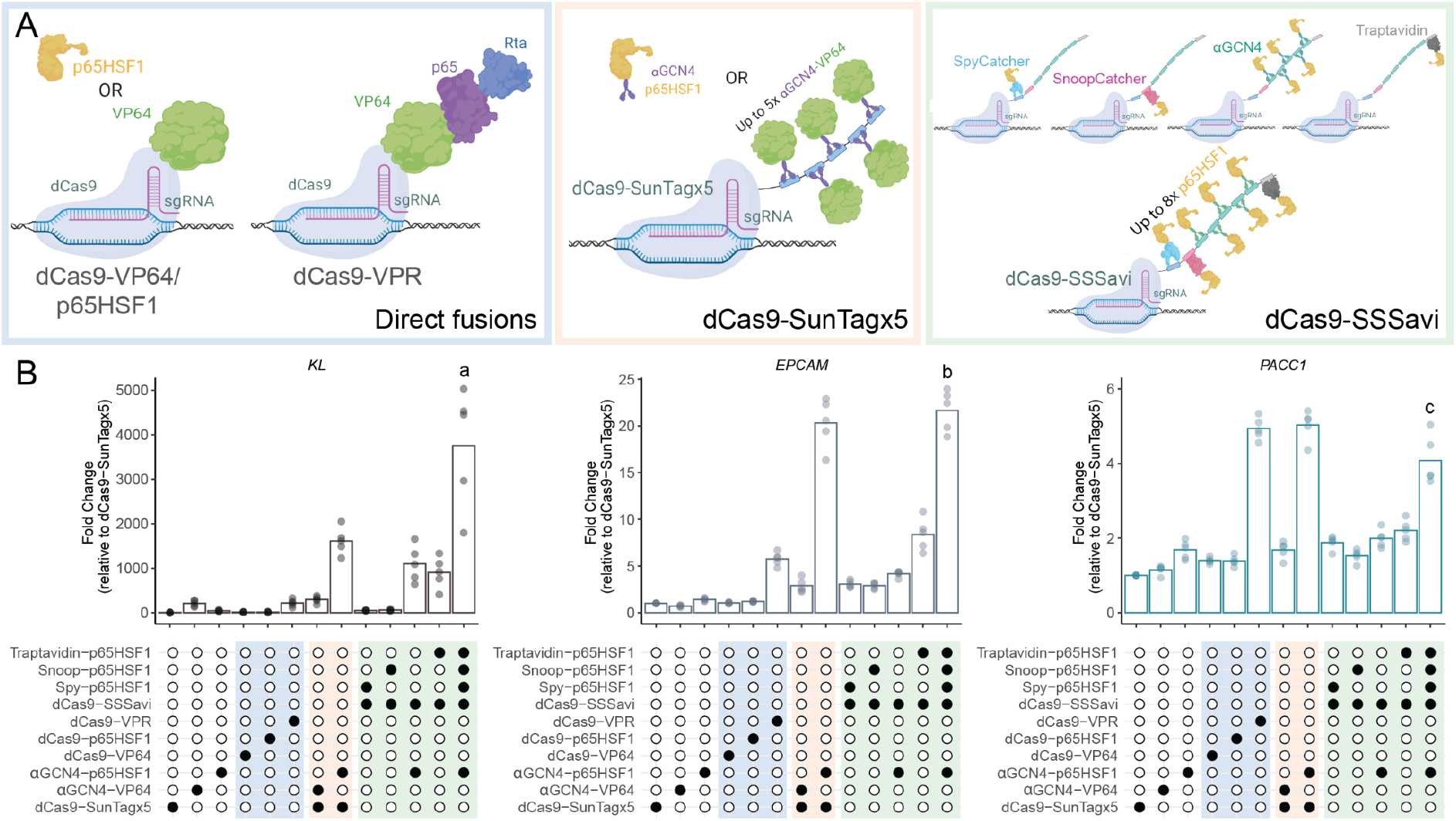
Comparison of SSSavi to other activation systems. **(A)** A schematic of the three activation systems tested: direct fusions (dCas9 fused to VP64, p65HSF1, or VPR), dCas9-SunTagx5 (recruiting VP64 or p65HSF1), or dCas9-SSSavi recruiting p65HSF1 fused to one of four catchers (SpyCatcher, SnoopCatcher, *α*GCN4, or Traptavidin). **(B)** RT-qPCR quantification of target gene transcriptional upregulation with different activation systems: direct fusions (blue), dCas9-SunTagx5 (orange), and dCas9-SSSavi (green). HEK293T 6x sgRNA stable cells (H6G) were used for transfection (n = 5 biological replicates, bar graphs indicate sample mean, fold change calculated based on dCas9-SunTagx5 only transfected cells, sorted for BFP and GFP). Bars with different letters indicate a significant difference as calculated by independent sample t-tests when comparing the SSSavi system with a full complement of catchers to the two strongest activation platforms tested (dCas9-VPR or dCas9-SunTagx5 while recruiting p65HSF1). a: significantly different to both platforms (*p* = 4.8×10^−6^ and 0.01, respectively), b: significantly different to VPR (*p =* 1.1×10^−7^), c: not significantly different to both systems (*p* = 0.04 and 0.03).

### Order of effector recruitment alters efficacy of transcriptional regulation

Having established the efficacy of the dCas9-SSSavi platform for strong and specific target gene activation, we next sought to characterise how well the SSSavi system performs for target gene transcriptional down regulation. To this end, the dCas9-SSSavi construct, along with the *E. coli* biotin holoenzyme synthetase, BirA, were stably integrated into the 6x sgRNA HEK293T stable line (H6G), referred to as HSB6G cells (HEK293T, dCas9-SSSavi, BirA, 6x sgRNAs). HSB6G cells were transiently transfected with one of three well-established repressive effectors, DNMT3A (D3A), KRAB (K), or EZH2 (E), linked to each of the four individual SSSavi catchers. Each catcher-effector was assessed independently for their repressive capacity, as well as co-transfection of all four catchers together with the same effector. Four of the most abundantly expressed target genes (*PACC1, B2M, RBM3*, and *HINT1*) were measured to assess reduction in their mRNA levels. To control for possible confounding effects of PEI-based transfection on the transcriptome, log_2_ fold change values for the different SSSavi constructs were calculated compared to transfection of an *α*GCN4-mCherry non-catalytic control. *α*GCN4-mCherry was chosen to account for potential non-catalytic inhibitory effects such as preventing physiological interactions of transcriptional regulators from binding to the promoter region. Consequently, we deemed targeted effector specific repression was only achieved if expression levels exceeded the baseline repression levels induced by mCherry alone. Overall, testing each effector independently revealed no significant repression by KRAB, while EZH2 induced significant downregulation of *PACC1* when recruited by *α*GCN4, and DNMT3A significantly reduced *HINT1* transcript abundance when recruited by Snoop or when fused to all four catchers simultaneously (**Figure 4A**).

**Figure 4.**
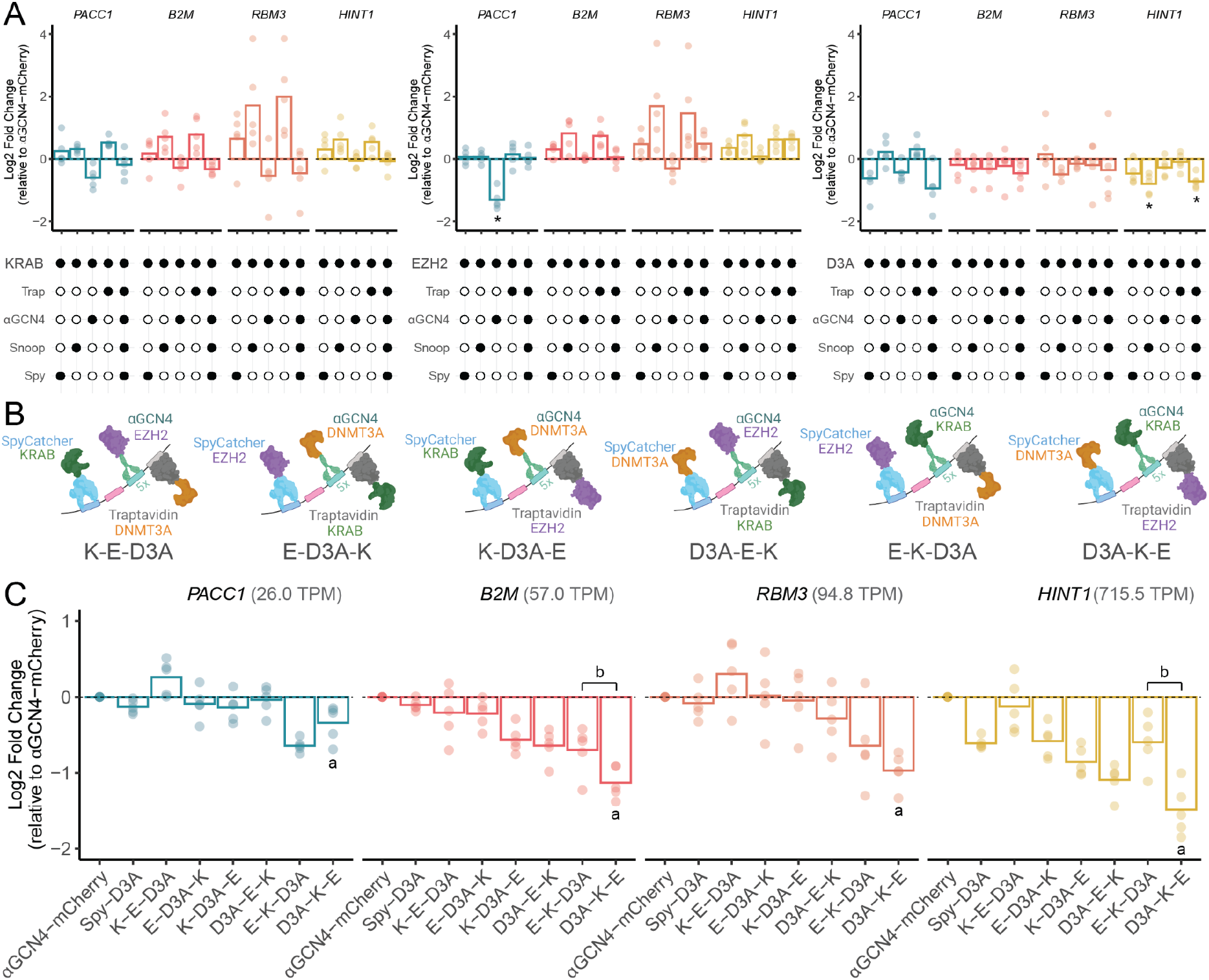
Recruitment of DNMT3A, KRAB, and EZH2 to induce targeted transcriptional repression via the SSSavi system. RT-qPCR quantification of target gene transcriptional repression induced by transient transfection of SSSavi plasmids in HSB6G (dCas9-SSSavi and 6x sgRNA) stable cells. **(A)** Catchers were tested individually or in four-way combinations for each of three effectors: DNMT3A (D3A), EZH2 (E), and/or KRAB (K) fused to either SpyCatcher, SnoopCatcher, *α*GCN4, or Traptavidin, across four different target genes (*PACC1, B2M, RBM3*, and *HINT1*). Fold change in expression was calculated compared to transfection with *α*GCN4-mCherry alone (baseline control, black dotted lines). *Independent sample t-tests with Benjamini-Hochberg correction comparing effector to *α*GCN4-mCherry, where *p*-values < 0.05. **(B)** A schematic of the six possible three-way combinations indicating the order in which effectors (D3A, K, and E) were recruited onto the SSSavi docking array due to their differential fusion to SpyCatcher, *α*GCN4, or Traptavidin. **(C)** RT-qPCR quantification of target gene transcript abundance following transient transfection with all possible three-way combinations of spatial recruitment D3A, K, and E on the SSSavi docking array. Two control samples were also included: *α*GCN4-mCherry as the non-catalytic baseline control (black dotted line) and SpyCatcher fused to D3A as a single epimodifier positive control. Bars with different letters indicate a significant difference as calculated by independent sample t-tests. a: compares D3A-K-E to aGCN4-mCherry, and b: compares D3A-K-E to E-K-D3A (*p*-values < 0.05). **(A**,**C)** Bar graphs indicate sample mean (n = 5, biological replicates, log_2_ fold change calculated compared to *α*GCN4-mCherry control samples, sorted for GFP).

Given these results, we sought to determine if combining these effectors might improve the level of observed downregulation. This hypothesis was underpinned by previous studies that demonstrated that combining DNMT3A with either KRAB or EZH2 led to persistent epigenetically induced gene silencing [9,21]. The key advantage of the SSSavi system is the ability to control the number and the order of effectors that bind to the docking array. Thus, we tested whether combinatorial recruitment of three different repressive domains would improve the level of gene silencing, and whether the degree of silencing was dependent on the linear spatial order in which the effectors were recruited onto the SSSavi docking array. This was done by testing all three-way combinations of D3A, K, and E, with the spatial order of recruitment on the SSSavi dock being SpyCatcher, *α*GCN4 and Traptavidin. This revealed two key findings: first, combinatorial recruitment resulted in amplified transcriptional repression; and second, the arrangement in which the effectors are recruited significantly affects the strength of the repression observed (**Figure 4.B**). Specifically, the SpyCatcher-D3A, *α*GCN4-KRAB, and Traptavidin-EZH2 combination (denoted as D3A-K-E) showed greater repression than all other permutations at three of the four target genes, with *p*-values provided in **Supplementary Table S4** when compared to either the *α*GCN4-mCherry control, or to the next strongest combination, E-K-D3A). The only exception was the E-K-D3A arrangement, which showed stronger repression (log_2_ fold change of −0.64) than D3A-K-E at *PACC1*. When the same three effectors were recruited as

K-E-D3A, minimal repression was observed across all four genes. Furthermore, by combining the individual data (**Figure 4A**) with the combinatorial data (**Figure 4B**) and grouping across target genes to have sufficient sampling, we could perform an ANOVA to determine if an additive or synergistic effect was occuring. This analysis provided strong evidence that a synergistic effect was occurring for the two combinations that showed the greatest repression (D3A-K-E: *F* = 4.61, *p* = 0.017, E-K-D3A: *F* = 3.76, *p* = 0.033). That is, the transcriptional change induced by combining these three effectors in a predefined order was greater than that achieved by simply summing the effect of each individual effector alone. Overall, we were able to show that recruiting multiple different effectors amplifies the gene silencing effect achievable, as well as the importance of the linear spatial ordering in which these effectors are recruited.

### Downregulation of the liver cancer biomarker *EPCAM* by combinatorial editing

Having established the SSSavi system in HEK293T cells, we next sought to establish this editing platform for a more clinically relevant target. *EPCAM* was selected as it is a liver cancer biomarker found in invasive hepatocellular carcinoma (HCC) for which small interfering RNA (siRNA) [44–46] or chemotherapeutic agents such as doxorubicin [47] have previously been used to downregulate its expression. Focus on EPCAM is based on the finding that only a subset of HCC cells overexpress this cell surface marker, an indicator of cancer stem cells, which are associated with cancer proliferation and invasiveness [46]. Thus, by using siRNA or doxorubicin to target and reduce EPCAM expression, these studies have shown that they can attenuate HCC severity and growth. Based on this and as a further test of the combinatorial capacity and targeted ordering of the SSSavi system, we sought to determine if the same three-way effector combination recruited in HSB6G cells (D3A-K-E) could be utilised to downregulate *EPCAM* expression in HepG2 cells, a liver HCC line in which *EPCAM* is highly expressed (261.1 TPM, compared to 5.2 in HEK293T cells).

Similar to the prior experiments in HEK293T cells, we stably integrated the dCas9-SSSavi and BirA components into WT HepG2 cells, referred to as HepSB cells. These cells were then transiently co-transfected with the previously used 6x sgRNAs, as well as three-way combinations of the catchers linked to the repressive effectors KRAB (K), DNMT3A (D3A), and EZH2 (E). To simplify testing, only the weakest (K-E-D3A) and most effective (D3A-K-E) repressive combinations were examined, as established above in HSB6G cells (**Figure 4B**). As shown in **Figure 5A**, downregulation of three of the four measured target genes was achieved when recruiting D3A-K-E, compared to *α*GCN4-mCherry alone. Furthermore, a >10% reduction in EPCAM positive cells coincided with the downregulation of *EPCAM* mRNA expression for D3A-K-E transfected cells, as measured by flow cytometry of matched immunostained samples in comparison to cells transfected with *α*GCN4-mCherry (**Figure 5B**). Thus, the SSSavi system provides an additional tool for attenuating the levels of EPCAM mRNA and protein to a comparable extent as reported by doxorubicin treatment [47]. This result further highlights the SSSavi system’s unique ability to control the order of effector recruitment to effectively repress or activate target genes.

**Figure 5.**
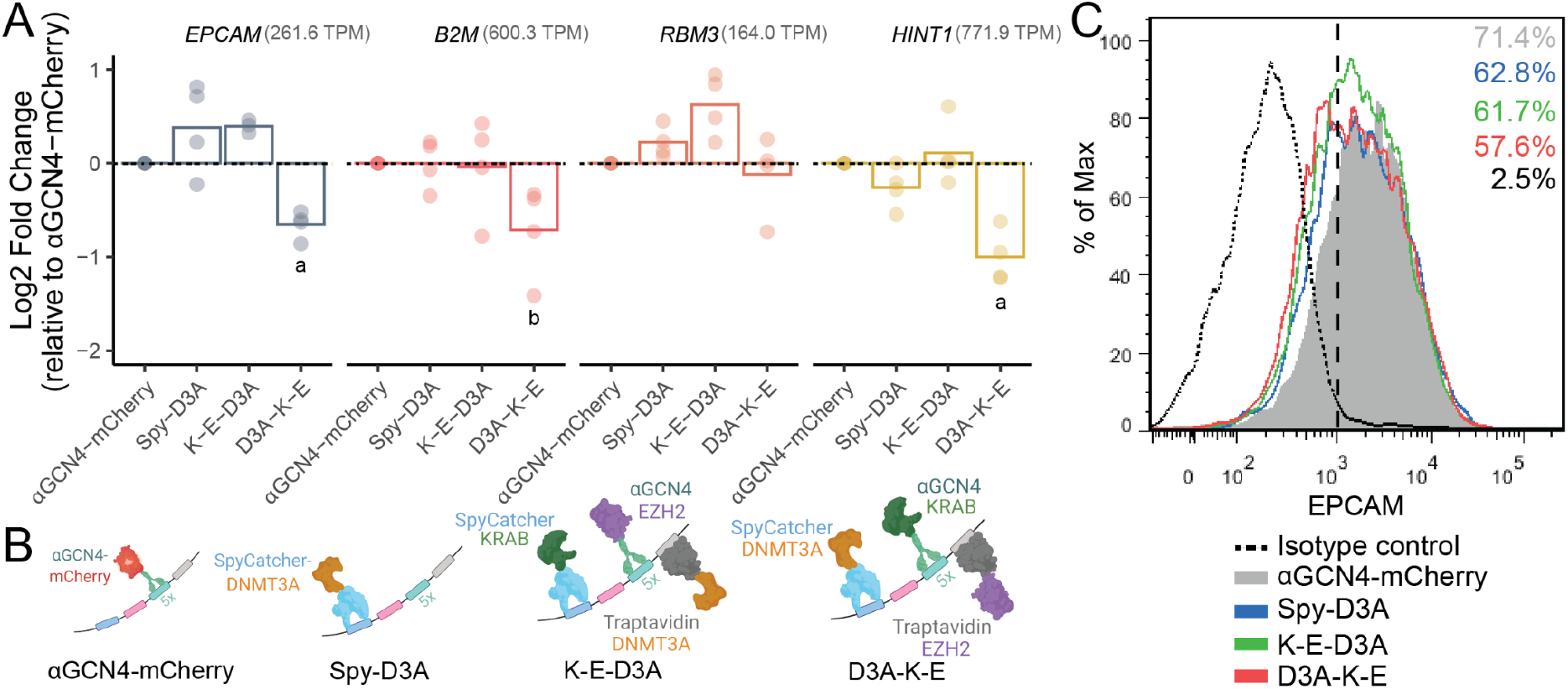
Downregulation of EPCAM mRNA and protein levels in HepG2 liver cancer cells by the SSSavi system. **(A)** RT-qPCR quantification of target gene transcriptional repression induced by transient transfection of SSSavi catcher plasmids in the HepSB (HepG2 cells with dCas9-SSSavi and BirA stable) cell line using DNMT3A (D3A), EZH2 (E), and KRAB (K), as well as a 6x sgRNA plasmid. Fold change in expression is calculated compared to transfection with *α*GCN4-mCherry alone (baseline control, black dotted lines). Bar graphs indicate sample mean with each letter indicating a significant difference as calculated by independent sample t-tests comparing D3A-K-E to *α*GCN4-mCherry (a: *p* = 0.003 (*KL*), 0.006 (*RBM3*), b: *p* = 0.06 (*B2M*)). n = 4, biological replicates, log_2_ fold change calculated compared to *α*GCN4-mCherry control samples, sorted for GFP (catchers) and mCherry (6x sgRNA). **(B)** A schematic of the different effectors recruited onto the SSSavi docking array, highlighting the different spatial ordering. **(C)** Flow cytometry analysis of HepSB cells immunostained with EPCAM antibody, comparing percentage of positive cells (vertical black dotted line) across four samples transfected with *α*GCN4-mCherry alone (grey), Spy-D3A alone (blue), K-E-D3A (green), or D3A-K-E (red) effector combinations. Duplicate experiments were performed.

## DISCUSSION

Here, we describe the design and validation of the novel SSSavi combinatorial recruitment platform that, when linked to dCas9, enables the targeting of up to four different effector domains to multiple target genes simultaneously to alter gene transcription. The four tag-catcher pairs demonstrate high binding specificity with excellent activation capabilities comparable to or exceeding currently existing CRISPR activation platforms, including SunTag and VPR. Furthermore, the SSSavi array enables potent transcriptional downregulation through the recruitment of multiple repressive domains, specifically DNMT3A, KRAB, and EZH2 in both HEK293T and HepG2 cells. This combinatorial variation underlines the importance of the spatial ordering or arrangement of epigenetic effectors recruited to a target locus.

Since the SSSavi contains five GCN4 peptides separated by linkers as part of the SunTag docking component, we initially hypothesised that the five binding domains may increase the recruitment of enzymatic effectors to induce stronger repression based by increasing the local presence of repressive enzyme domains such as DNMT3A or EZH2. To our surprise, the top two performing effector combinations (D3A-K-E and E-K-D3A) both have the non-catalytic KRAB repressive domain fused to *α*GCN4. Theoretically, this allowed for up to five copies of KRAB to be recruited to dCas9-SSSavi. Thus, *α*GCN4-KRAB appears to be playing a central role in the greater observed transcriptional downregulation, though the observed synergy upon the additional recruitment of DNMT3A and EZH2 (**Figure 4B**) far exceeds that achieved by either *α*GCN4-KRAB alone, or when recruiting KRAB fused to all four catchers (**Figure 4A**). Consequently, the magnitude of transcriptional repression achieved is dependent upon the combined recruitment of these three repressive domains in a predefined order, emphasising the advantage of SSSavi to elucidate this requirement.

Of note, order-dependent regulatory effects of epigenetic effectors directly fused to dCas9 have been observed previously [6,7]. However, the direct fusion design limits the versatility for testing alternative effector combinations. This is partially overcome by the modular SunTag platform, but due to its utilisation of a single epitope-antibody combination, the arrangement of different effectors cannot be predefined. Thus, the key strength of the SSSavi system as an editing tool is the ease of switching and testing multiple different domains while also providing the necessary control over stoichiometry and order of effector binding. Testing of additional protein-protein interaction domains in the future could further expand the docking array complexity, such as the recently reported coiled-coil dimer sets [31].

There are a wide array of future areas to which SSSavi could be applied. It provides the means to disentangle currently unclear or contradictory gene regulatory roles of a range of effectors found in different genomic contexts and cell types. An example is histone lysine methyltransferase G9A, an effector that has largely been associated with transcriptional repression but also suggested to have both activation and repression roles depending on interactor proteins and chromatin placement [51]. Another example is X chromosome inactivation (XCI), which involves the interaction of multiple epigenomic layers to achieve transcriptional repression, including via DNA methylation, histone modifications, and regulation of chromatin architecture [53]. Consequently, combinatorial epigenome editing may provide a means to recapitulate or reverse the epigenetic modifications that underpin XCI silencing in a locus-specific manner. Thus, the SSSavi recruitment platform may enable the precise alteration of particular combinatorial chromatin states using artificial effectors where the combinatorial and interconnected nature of the epigenome may have impeded alteration in the past [54,55].

Overall, our findings indicate that the SSSavi system is an effective transcriptional regulator with the novel capability of flexible, directed, combinatorial effector recruitment. Thus, SSSavi’s unique flexibility provides the means to more easily explore possible synergistic effectors via co-recruitment to specific loci to magnify their regulatory capacity and potentially alter chromatin contexts that have previously shown limited susceptibility to targeted epigenomic editing [56–58]. Additionally there is the possibility to functionally characterise the combinatorial effects of chromatin regulators on transcription and identify effectors combinations that elicit changes of different directionality, magnitude, and stability. Therefore, this work provides a useful new tool to explore the complex combinatorial and multilayered nature of the epigenome.

## DATA AVAILABILITY

All data analysed and used to generate all figures in this study are supplied in supplementary information below. Flow cytometry data for Figure 5C can be accessed at https://flowrepository.org/ Repository ID: FR-FCM-Z5GQ.

## ACKNOWLEDGEMENTS

Genomic data was generated at the Australian Cancer Research Foundation Centre for Advanced Cancer Genomics and Genomics WA. Flow cytometry and FACS data was obtained either through the Centre of Microscopy and Characterisation and Analysis (CMCA), the Telethon Kids Institute (TKI), or the Harry Perkins FACS Facility (Perkins). We would like to thank P. Blancafort for the kind donation of the HEK293T and VPR plasmid, P. Leedman for the HepG2 cells and J. Polo for the PiggyBac plasmids (piggyBac_pCAGG_Amp_gateway_Hygro and hyPBase). SSSavi images were created using BioRender.com.

## AUTHOR CONTRIBUTIONS

T.S., C.P. and R.L. conceived of the project and designed the experiments. T.S. and C.P. conducted the experiments. C.P. wrote the in-house plasmid-seq mapping pipeline. S.F., T.S and C.P. performed the statistical analysis. D.P. and J.P. conducted the sequencing. T.N. assisted with the FACS, immunostaining and RT-qPCR design. T.S. and J.L. performed the EPCAM immunostaining and flow analysis. T.S., C.P. and R.L. wrote the manuscript. All authors approved of and contributed to the final version of the manuscript.

## MATERIALS AND METHODS

### Cell culturing

HEK293T and HepG2 cells were grown in a humidified cell culture incubator at 37°C with 5% (v/v) CO_2_. HEK293T cells were grown in DMEM (Dulbecco’s Modified Eagle Medium) and the HepG2 cells in EMEM (Eagle’s Minimum Essential Medium). Both were supplemented with 10% (v/v) FBS (Moregate BioTech, #23301121), 1X (v/v) Glutamax (Life Technologies, #35050061) and 1% MEM Non-essential amino acid solution (ThermoFisher, #11140050).

### Cloning the SSSavi plasmids

#### dCas9-SSSavi

A dCas9 plasmid that contained one of each of the four tags, was constructed from the LLP457_pGK-dCas9-SunTag-BFP plasmid (Addgene, #100957) with each tag separated by a 22 amino acid linker [37] using an IDT gBlock to produce dCas9-Spy-Snoop-Sun-Avi-Tag-BFP. SpyTag and SnoopTag sequences were obtained from pET28a-SnoopTag-mEGFP-SpyTag (Addgene, #72325), while the AviTag sequence originated from AviTag-SpyCatcher plasmid (Addgene, #72326). This dCas9 plasmid was then used as the backbone into which four additional SunTag domains were inserted to clone dCas9-Spy-Snoop-**Sunx5**-AviTag-BFP with a 3xHA epitope tag at the N-terminus of dCas9 (dCas9-SSSavi-BFP, **Figure 1C.i**). This was achieved by linearizing the backbone with MluI and then performing Gibson assemby [59] with the four additional SunTags that were PCR amplified from pCAG-dCas9-5xPlat2AflD (Addgene, #82560).

#### Catcher-p65HSF1 (Spy, Snoop, αGCN4, Traptavidin)

The *α*GCN4-p65HSF1-CO (codon optimised: CO) plasmid (**Figure 1C.ii**) was used as the template for all catcher-effector constructs. It was synthesised by PCR amplifying the *α*GCN4 backbone from LLP252 pEF1a-NLS-scFvGCN4-DNMT3a (Addgene, #100941) and the first 639 bp of the 5’ end of p65HSF1 from pAC1393-pmax-NLSPUFa_p65HSF1 (Addgene, #71897). These were then Gibson assembled along with an IDT gBlock containing the last 300 bp of p65HSF1 that had been codon optimised for expression level detection distinct from endogenous mRNA. The Spy and Snoop catcher domains were cloned from the pET28a-SpyCatcher-SnoopCatcher plasmid (Addgene, #72324) while the Traptavidin domain came from pET21a-Core-Traptavidin (Addgene, #26054). Each catcher was individually Gibson assembled into the *α*GCN4-p65HSF1-CO backbone by replacing the *α*GCN4, sfGFP and GB1 domains (**Figure 1C.iii**). A pSV40-GFP cassette was added for FACS purposes using Gibson assembly by PCR amplifying the GFP domain from an in-house pEF1a-GFP-Puro plasmid and the pSV40 promoter from an in-house dCas9-SunTag lentiviral plasmid (pLenti-pSV40-dCas9-SunTag-P2A-BFP-WPRE). Each Traptavidin construct was co-transfected with the E.coli biotin holoenzyme synthetase, pEF1a-BirA-V5-neo (Addgene, #100548) in order to add biotin to the lysine residue on the AviTag [60].

#### Repressive catcher-effectors

The three repressive domains examined (DNMT3A, KRAB, and EZH2) were ordered as codon optimised gBlocks from IDT and inserted into the respective catcher plasmids using NotI and HpaI. The control sample, *α*GCN4-mCherry, was cloned by PCR amplifying both the mCherry from LLP469 pEF1a-mCherry-EMPTY-gRNA (Addgene, #100958) and the backbone of *α*GCN4-p65HSF1-CO, followed by Gibson assembly, replacing the p65HSF1 domain with mCherry.

#### dCas9-SSSavi piggyBac plasmid

To create the construct needed for establishing a SSSavi stable expression line (HSB6G), we used Gateway cloning (Invitrogen) to insert dCas9-SSSavi and BirA into a piggyBac plasmid [61]. The dCas9-SSSavi was PCR amplified to include attB sites and then inserted into pDONR221 (Invitrogen) using BP clonase II (Thermo Fisher, #11789020) to create the pENTRY221_dCas9-SSSavi plasmid. This was then linearized with EcoRV and NotI into which BirA was inserted to clone pENTRY221_dCas9-SSSavi_BirA. The dCas9-SSSavi_BirA cassette was subsequently inserted into PB_CAH (piggyBac_pCAGG_Amp_gateway_Hygro) using LR Clonase II (Thermo Fisher, #11791020) to create the plasmid PB_dCas9-SSSavi_BirA.

#### Single use plasmids for characterising the SSSavi system

The plasmid sequences for the single use plasmids for characterising the SSSavi system including the dCas9 constructs tested for binding specificity (**Figure 2**), the direct fusion plasmids dCas9-VP64, dCas9-p65HSF1, dCas9-VPR (**Fig 3**), the switched dCas9-catcher and tag-effector fusions (**SuppFig S1**), the plasmids used to assess catcher protein stability (**SuppFig S3**) and the dCas9 SSSavi platform with 1xSunTag (**SuppFig S4**) and are detailed in **Supplementary Information S1**.

#### sgRNA plasmids

Six target genes were selected and their transcript per million (TPM) values in HEK293T cells based on in-house RNA-seq data were: *KL*, 0.01 TMP (*Klotho*, [62], *EPCAM*, 5.2 TPM (*epithelial cellular adhesion molecule*, [63,64], *PACC1*, 26.0 TPM (*proton activated chloride channel 1*, aka *TMEM206*, [40], *B2M*, 57.0 TPM (*beta-2-microglobulin*, [9], *RBM3*, 94.8 TPM (*RNA binding motif protein 3*), and *HINT1*, 715.5 TPM (*histidine triad nucleotide binding protein 1*). These sgRNAs were first cloned individually into the backbone LLP469 pEF1a-mCherry-EMPTY-gRNA (Addgene, #100958) following the protocol by Mali et al. [65]. The final 6x sgRNA plasmid was synthesised using Golden Gate cloning [66] and then transferred into pLenti-Puro (Addgene, #39481) using restriction ligation with SpeI and KpnI to create pLenti-6xHRBKET-sgRNA (6x sgRNA). All sgRNA sequences are reported in **Supplementary Table S1**.

#### Plasmid sequence verification

Plasmids were sequence verified using an in-house Tn5 tagmentation library preparation protocol based on work by [67–69]. Libraries were run on an Illumina MiSeq with paired-end reads, 150 bp fragments and the resulting FASTQ files were assembled using an in-house plasmid-mapper pipeline that incorporates *SPAdes* de novo assembly [70], unicycler and Bowtie2 alignment [71] with a reference FASTA file and the resulting contigs were sequence verified using IGV [72] and SnapGene software (GSL Biotech). A list of all plasmids is reported in **Supplementary Excel File S1** detailing those that have been deposited at Addgene and those that have the plasmid sequence reported in **Supplementary Information S1-20**.

### Cell transfections

Transfections were carried out using a 1:3 ratio of DNA to polyethylenimine, branched MW 25,000 Da (PEI, Sigma-Aldrich, 408727). Briefly, cells were seeded in either 24-well (In Vitro Technologies, #FAL353047) or 6-well culture plates (In Vitro Technologies, #FAL353046) at 40,000 or 200,000 cells, respectively. Cells were transfected with an equal ratio of plasmids 24 h post-seeding, with either 500 ng total DNA per 24-well, or 2.5 µg per 6-well plate and 1.5 µL or 7.5 µL of PEI (1 mg/mL), and 50 µL or 250 µL OptiMEM. Cell media was replaced after 3.5 h to remove PEI and minimise cell toxicity. Subsequently, cells were harvested 72 h post transfection (ptf) for FACS and RT-qPCR.

### Stable cell lines

Experiments were performed in either WT HEK293T cells or one of three stable cell lines. These included (1) <u>H</u>EK293T cells with the <u>6</u>x s<u>g</u>RNA stably integrated (H6G), which were used to produce the subsequent transgenic lines: (2) <u>H</u>EK293T cells expressing dCas9-<u>S</u>SSavi, <u>B</u>irA and <u>6</u>x s<u>g</u>RNA (HSB6G), and finally, (3) <u>Hep</u>G2 cells stably expressing dCas9-<u>S</u>SSavi and <u>B</u>irA (HepSB). The creation of the H6G line was achieved using third generation lentiviral transduction, followed by clonal selection of pLenti-6xHRBKET-sgRNA. The HSB6G and HepSB SSSavi lines utilised the piggyBac system for stable integration and expression [61]. Specifically, these cells were established by transient co-transfection of H6G and HepG2 WT cells, respectively, with the PB_dCas9-SSSavi-BirA plasmid and hyPBase, followed by hygromycin selection (250 µg/mL) for 2 weeks. Integration and stable expression of dCas9 was confirmed using immunostaining followed by flow cytometry as described below.

### Verifying stable integration of dCas9 using immunostaining

HEK293T cells were collected and fixed in 2% paraformaldehyde before being incubated in blocking buffer (1x PBS, 5% Goat serum, 0.3% Triton-X100) for 1 hr at room temperature. The blocking buffer was subsequently removed and cells were incubated for an additional hour with 1:100 diluted primary antibody (*α*HA antibody, Biolegend, #901502) in antibody dilution buffer (1x PBS, 2.5% Goat serum 0.2%, Triton-X100). Cells were washed three times in antibody dilution buffer and then incubated in the dark for 30 min with 1:1000 diluted secondary antibody (Goat anti-mouse AF488, Biotium, #12C0403). Cells were again washed three times before being resuspended in sort buffer (1x PBS, 2.5% FBS, 5 mM EDTA) and analysed using flow cytometry to determine the percentage of AF488/GFP positive cells.

### EPCAM immunostaining in HepG2 cells

Transfected HepSB cells were collected 72 hrs ptf, with 25% of the cells used for immunostaining and the remainder being isolated by FACS for RT-qPCR analysis. Cells utilised for immunostaining were incubated in blocking buffer (1x PBS, 1% BSA, 1 mM EDTA) for 30 min at room temperature. Cells were subsequently incubated for an additional hour with 1:10 diluted primary antibody (PE/Cy7 anti-human CD326 (EPCAM) Antibody, Biolegend, #369815) in antibody dilution buffer (1x PBS, 1% BSA, 1 mM EDTA). Cells were then washed four times in antibody dilution buffer before being resuspended in sort buffer (1x PBS, 2.5% FBS, 5 mM EDTA) and >5,000 singlet, GFP positive cells were analysed per sample using flow cytometry to determine the percentage of PE-Cy7/EPCAM positive cells.

### FACS and flow cytometry

Flow cytometry was performed using the BD FACSCanto to determine the percentage of AF488/GFP positive cells as an indicator of dCas9 stably expressing cells after immunostaining. A BD FACSAria II was used for all HepG2 flow cytometry and FACS experiments. For HEK293T samples that were isolated by FACS using a BD FACSAria III, a BD FACSMelody, or a BD FACSAria II. Cell samples isolated by FACS were centrifuged at 300 xg for 5 min and the supernatant removed in preparation for RNA extraction and RT-qPCR.

### RNA extraction

An in-house bead based RNA extraction protocol was used on <100,000 cells isolated by FACS. Briefly, pelleted cells were resuspended in 50 µL of 1x PBS and an equal volume of cell homogenization buffer (6 M Guanidine Thiocyanate (Astral Scientific, #BIOGB0244), 50 mM Tris HCl, pH 8.0 (Thermo Fisher, #AM9856), 2% (w/v) Sarkosyl (Sigma, #L9150), 20 mM EDTA (Life Tech, #AM9260G) in nuclease free water (Thermo Fisher, #AM9937) containing 1:100 β-mercaptoethanol (Astral Scientific, #AM0482). The cell suspensions were lysed in 300 µL of RNA lysis buffer (6 M Guanidine Thiocyanate and 100 mM Tris Base, pH 7.3-7.7, in nuclease free water) and vortexed. 0.7X volume of isopropanol was then added to each sample and briefly vortexed. To this mixture, 10 µL of Mag-Binding beads (Zymo Research, #D4100-2-24) was added and the sample was incubated for 3 min at room temperature, shaking at 1500 rpm. The supernatant was then removed from each sample using a Dynamag (Thermo Fisher, #12321D) and washed with 500 µL AW1 buffer (6 M Guanidine Thiocyanate, 95% EtOH, pH 5) followed by three washes with 80% EtOH. Beads were then dried before adding 50 µL DNaseI mix (3 U of DNaseI in 1X DNAse buffer (NEB, #M0303L) and incubated at 37 °C for 10 min, shaking at 1300 rpm. To this, 300 µL of AW1 buffer was added and incubated at room temperature for 3 min, shaking 1500 rpm. Samples were then returned to the magnetic plate and the beads subsequently washed twice in 80% EtOH. After the final wash, the beads were dried before the RNA was eluted in 30 µL nuclease free water. RNA samples were then used for cDNA synthesis and qPCR.

### RT-qPCR

Up to 900 ng of RNA was used per sample to synthesise cDNA using the SensiFAST cDNA Synthesis kit (Bioline, #BIO-65054) following the manufacturer’s protocol. qPCR primers (**Supplementary Table S2**) were designed using Primer 3 (http://primer3.ut.ee/) for three housekeeping (HK) controls, *GAPDH, RPS18*, and *HSPC3* and for target genes *KL, EPCAM, PACC1, B2M, RBM3*, and *HINT1*. qPCR was performed using 5 µL of 2X Luna Universal qPCR Master Mix (NEB, #M3003E), 2 µL of 10 µM primer pair mix and 2 µL of 5 ng/µL cDNA per reaction. Samples were then run on the Applied Biosystems ViiA 7 instrument (ThermoFisher, #4453545) using the following program: 95°C for 3 min, followed by 40 cycles at 95°C for 10 sec, 63°C for 20 sec and then 72°C for 5 sec.

## Data analysis

Gene expression levels were normalized to the geometric mean of the three HK control genes with ΔΔCt calculated by comparing to control samples as stated on the y-axis of figures. For measuring repression, values were compared to transfection of an *α*GCN4-mCherry non-catalytic control. Thus any downregulation seen was not solely due to catcher binding to the SSSavi platform but a result of effector induced repression. Additionally, *KL* expression level was set to an arbitrary baseline C_T_ value of 35, as a cutoff of no transcripts present. Independent sample t-tests with Benjamini-Hochberg multiple test corrections were used to calculate statistical significance where stated. Each figure details the number of replicates for each experiment.

## Supplementary Figures

**Supplementary Figure S1.**
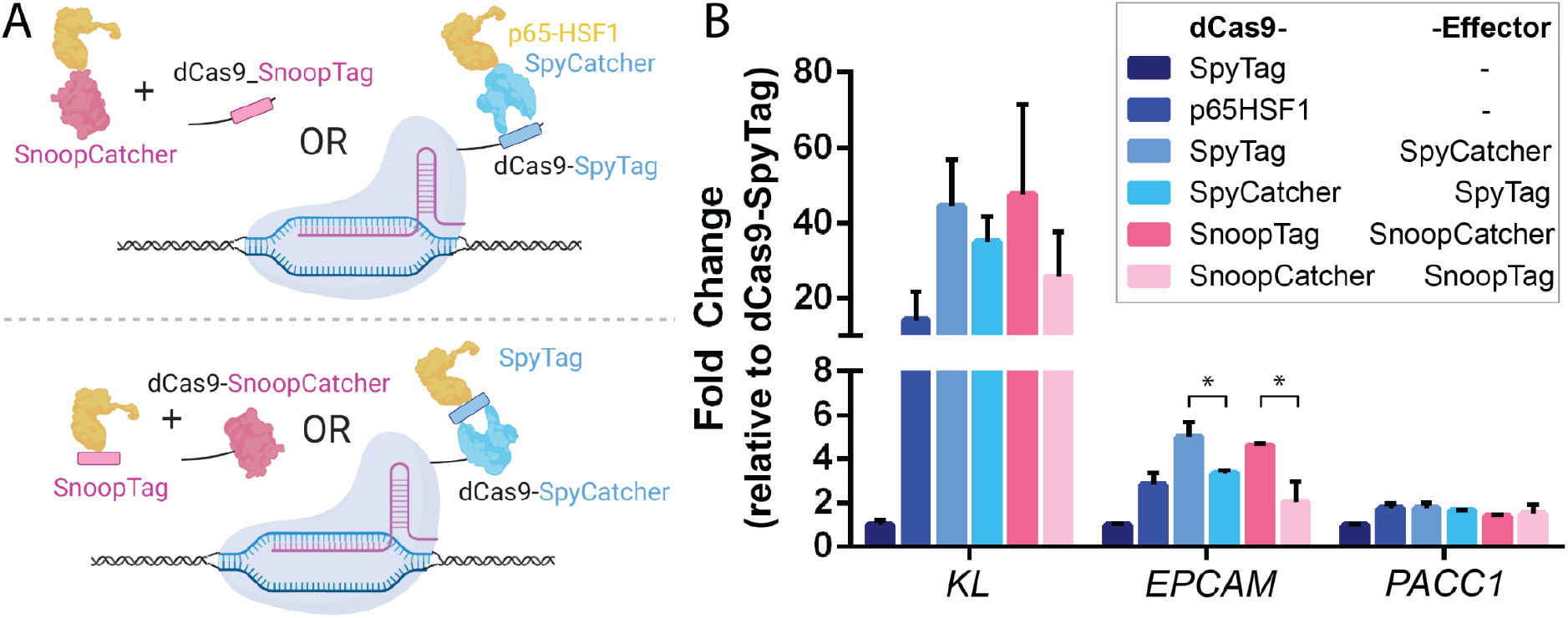
Comparison of fusing the tag or catcher domains to either dCas9 or effector. **(A)** A schematic of the dCas9-tag and catcher-effector fusions (top left) compared to the dCas9-catcher and tag-effector fusions (bottom left). **(B)** Fold change in the transcript abundance of three target genes following transfection of the different tag/catcher fusions, relative to transfection of the dCas9-SpyTag only construct. H6G cells (HEK293T cells stably expressing 6x sgRNAs) were used (n = 3, biological replicates, fold change calculated based on dCas9-SpyTag only transfected cells, sorted for BFP (dCas9) and GFP (catchers), mean ± SD, statistical significance determined using independent sample t-tests, \**p* < 0.05).

**Supplementary Figure S2.**
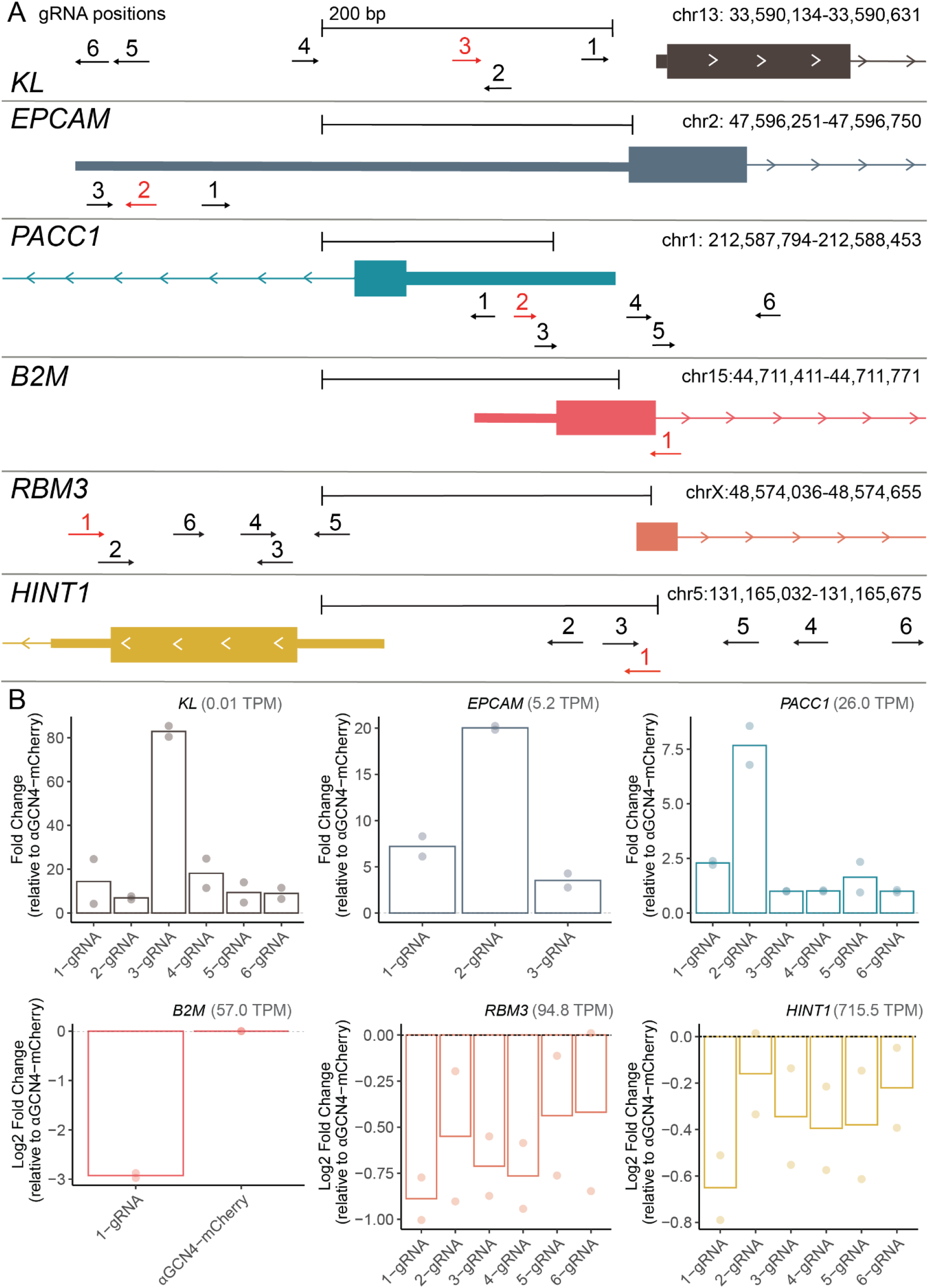
Identification of effective sgRNAs to use for activation/repression of *KL, EPCAM, PACC1, B2M, RBM3*, and *HINT1* target genes. **(A)** Positional information for up to six sgRNAs relative to the target gene promoter. The best performing sgRNA (red) was selected for stable integration in HEK293T cells based on **(B)** RT-qPCR quantitation of target gene transcript abundance. HEK293T cells were transiently transfected with dCas9-SSSavi and an individual sgRNA, along with either *α*GCN4-p65HSF1 for testing activation of the lowly expressed *KL, EPCAM*, and *PACC1*, or *α*GCN4-KRAB for testing repression of the highly expressed *B2M, RBM3*, and *HINT1*. Fold change in expression was calculated compared to transfection with *α*GCN4-mCherry alone (baseline control, n = 2, biological replicates, unsorted cells).

**Supplementary Figure S3.**
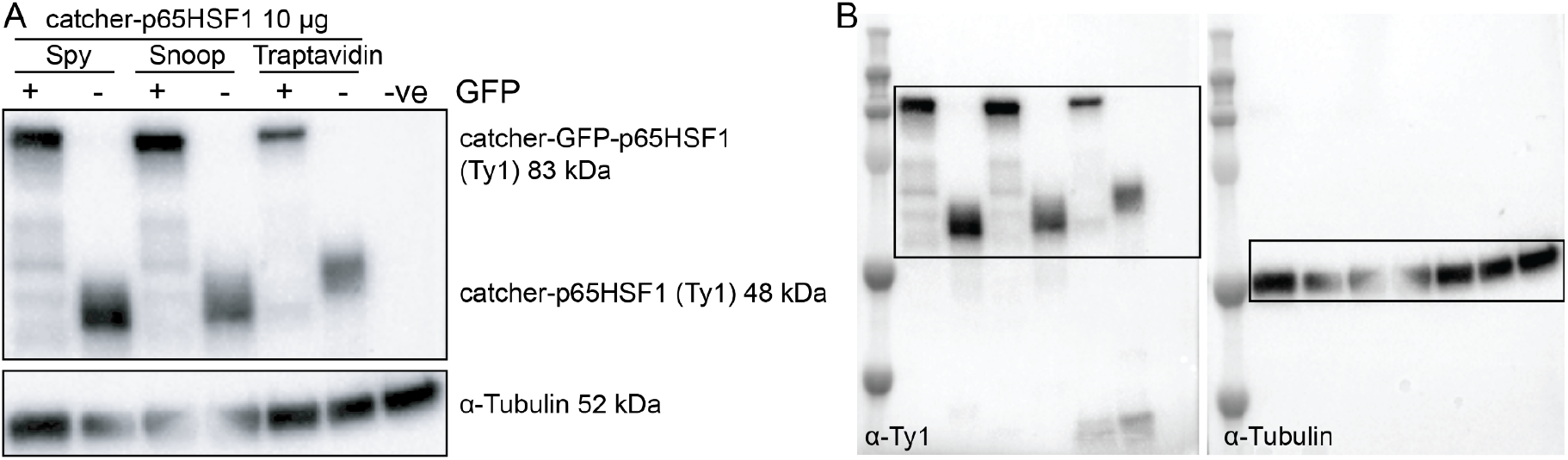
Protein expression and stability of different SSSavi constructs. **(A)** Western blot of catcher-effector proteins (anti-Ty1) per 10 µg of total protein from cell lysate. The blot indicates WT HEK293T cells transiently transfected with catchers fused to p65HSF1, either with (83 kDa) or without (48 kDa) internal GFP domains. -ve indicates untransfected HEK293T WT cells. Loading control: alpha-Tubulin. **(B)** Full unannotated western blots of **(A)**.

**Supplementary Figure S4.**
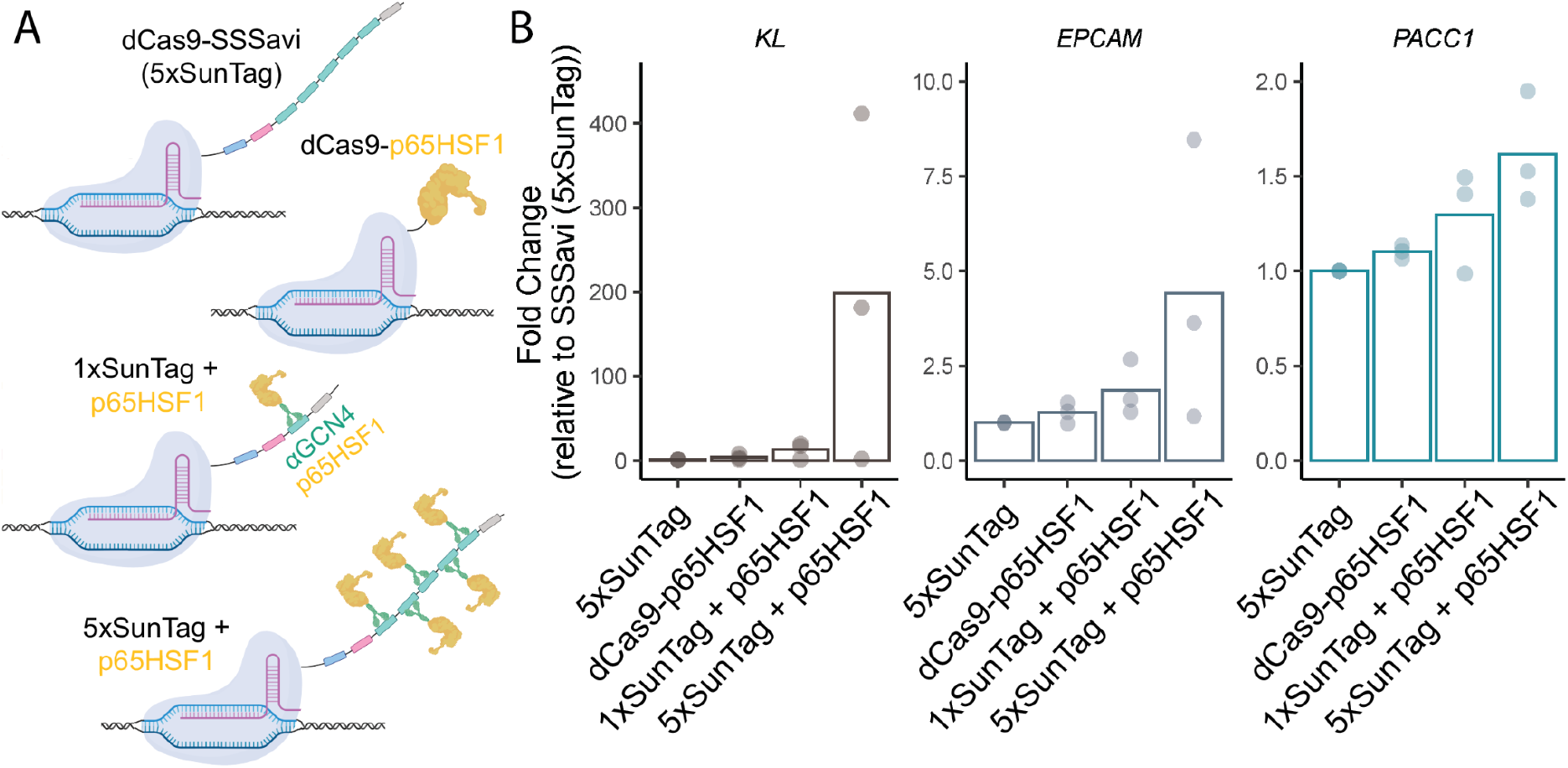
Comparison of 1x and 5x SunTag arrays for target gene activation when recruiting *α*GCN4-p65HSF1. **(A)** Schematic of the dCas9-SSSavi platforms with 1x or 5x GCN4 peptide repeats in the SunTag domain recruiting *α*GCN4-p65HSF1, with the direct fusion plasmid, dCas9-p65HSF1, as a positive control. **(B)** RT-qPCR quantification of the fold change in the transcript abundance of 3 target genes following transfection of H6G cells with dCas9-SSSavi (5xSunTag) alone (control), dCas9-p65HSF1, or dCas9-SSSavi containing 1x or 5x GCN4 repeats with *α*GCN4-p65HSF1. n = 3, biological replicates, fold change calculated based on dCas9-SSSavi (5xSunTag variant) transfected cells, BFP and GFP double-positive sorted cells.

## Supplementary Tables

**Supplementary Table S1.**
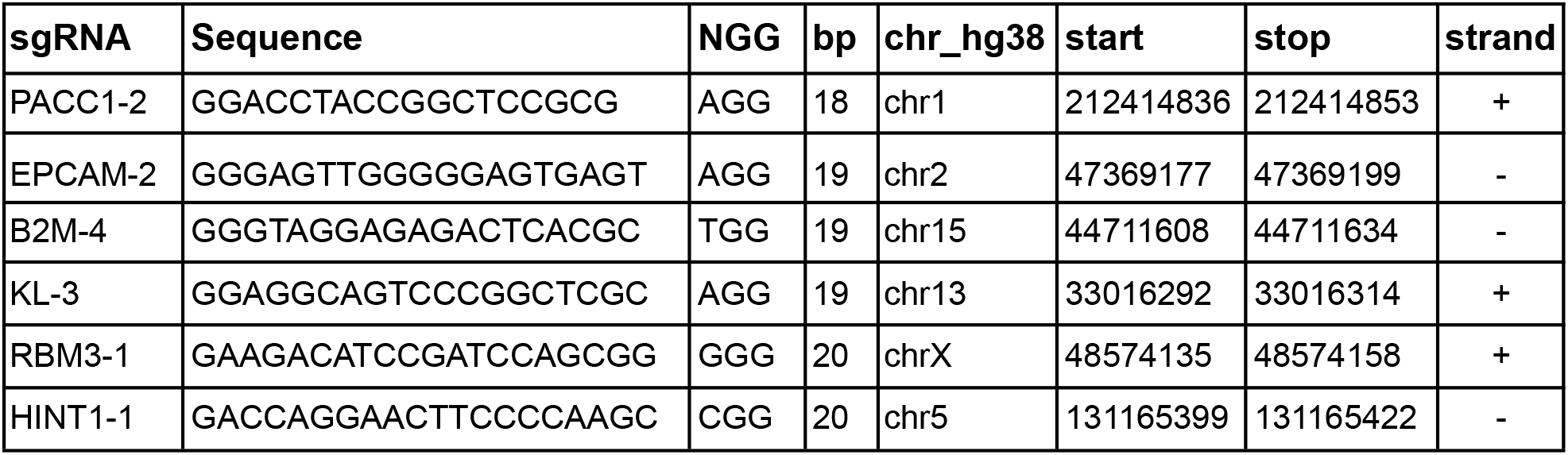
Guide RNA target sequence information for six target genes.

**Supplementary Table S2.**
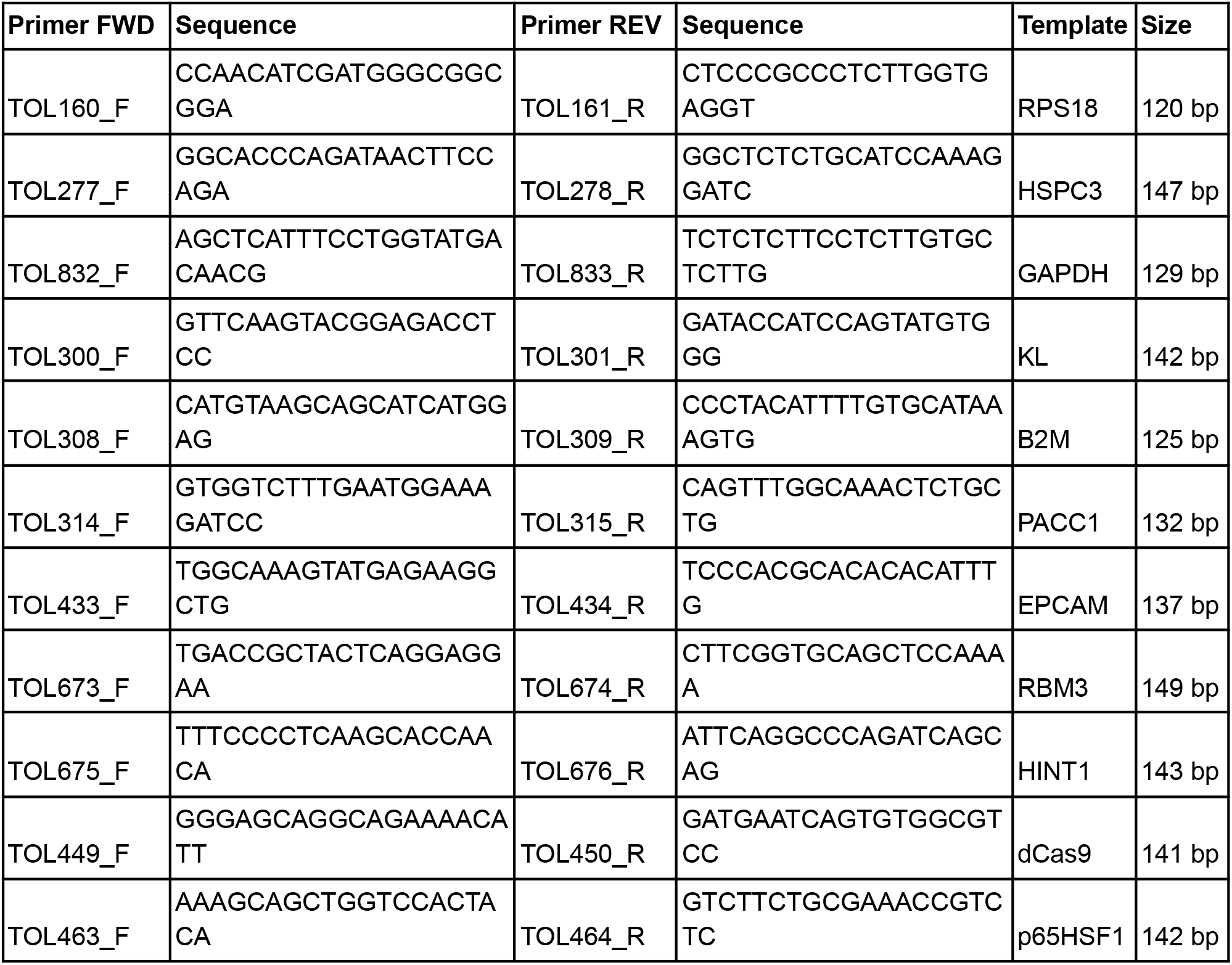
qRT-PCR primer sequences for three housekeepers (*RPS18, HSPC3, GAPDH*) and six target genes (*KL, EPCAM, PACC1, B2M, RBM3, HINT1*), as well as dCas9 and p65HSF1.

**Supplementary Table S3.** Independent sample t-tests with *p*-values comparing single or multiple catcher combinations for transcriptional activation associated with **Figure 1D** (n = 2). Values are provided for each of three target genes comparing dCas9-SSSavi only to single catchers (0 vs 1), single catchers to pairwise combinations (1 vs 2), pairwise to three-way combinations (2 vs 3), and three-way to four-way combinations (3 vs 4).

|  |  | T-test comparison |  |  |  |
| --- | --- | --- | --- | --- | --- |
| Gene |  | 0 vs 1 | 1 vs 2 | 2 vs 3 | 3 vs 4 |
| ***KL*** | **t-test** | 15.68 | 5.68 | 3.45 | 4.99 |
|  | ***p*-value** | 1.04x10<sup>-6</sup> | 2.94x10<sup>-5</sup> | 0.003 | 0.001 |
| ***EPCAM*** | **t-test** | 8.41 | 4.1 | 3.46 | 2.22 |
|  | ***p*-value** | 6.62x10<sup>-5</sup> | 0.001 | 0.004 | 0.059 |
| ***PACC1*** | **t-test** | 4.94 | 3.22 | 2.54 | 2.09 |
|  | ***p*-value** | 0.001 | 0.006 | 0.023 | 0.094 |

**Supplementary Table S4.** Independent sample t-tests with *p*-values comparing the strongest (D3A-K-E) and weakest (K-E-D3A) repressive three-way combinations to the non-catalytic control (αGCN4-mCherry), as well as comparing the two strongest combinations (E-K-D3A and D3A-K-E) across four different target genes as depicted in **Figure 4B** (n = 5).

| Gene | αGCN4-mCherry vs D3A-K-E |  | αGCN4-mCherry vs K-E-D3A |  | E-K-D3A vs D3A-K-E |  |
| --- | --- | --- | --- | --- | --- | --- |
|  | t-test | *p*-value | t-test | *p*-value | t-test | *p*-value |
| ***PACC1*** | -3.21 | 0.032 | 2.55 | 0.063 | 2.65 | 0.044 |
| ***B2M*** | -11.97 | 0.0003 | -1.32 | 0.257 | -2.53 | 0.039 |
| ***RBM3*** | -9.42 | 0.0007 | 1.59 | 0.186 | -1.26 | 0.260 |
| ***HINT1*** | -9.85 | 0.0006 | -0.78 | 0.478 | -4.17 | 0.003 |

## Supplementary Information

### Supp Info S1. SSP073_dCas9-VP64

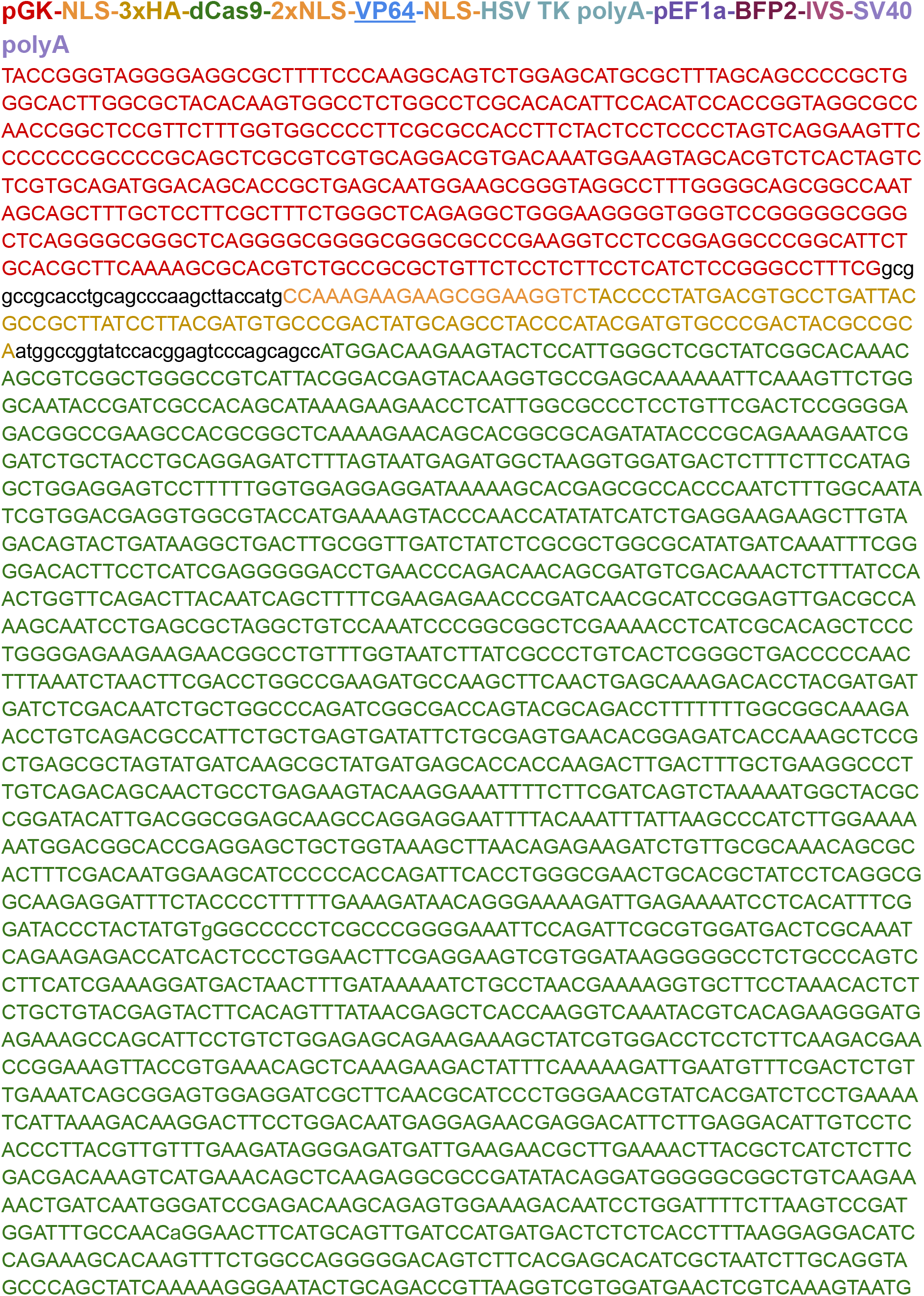
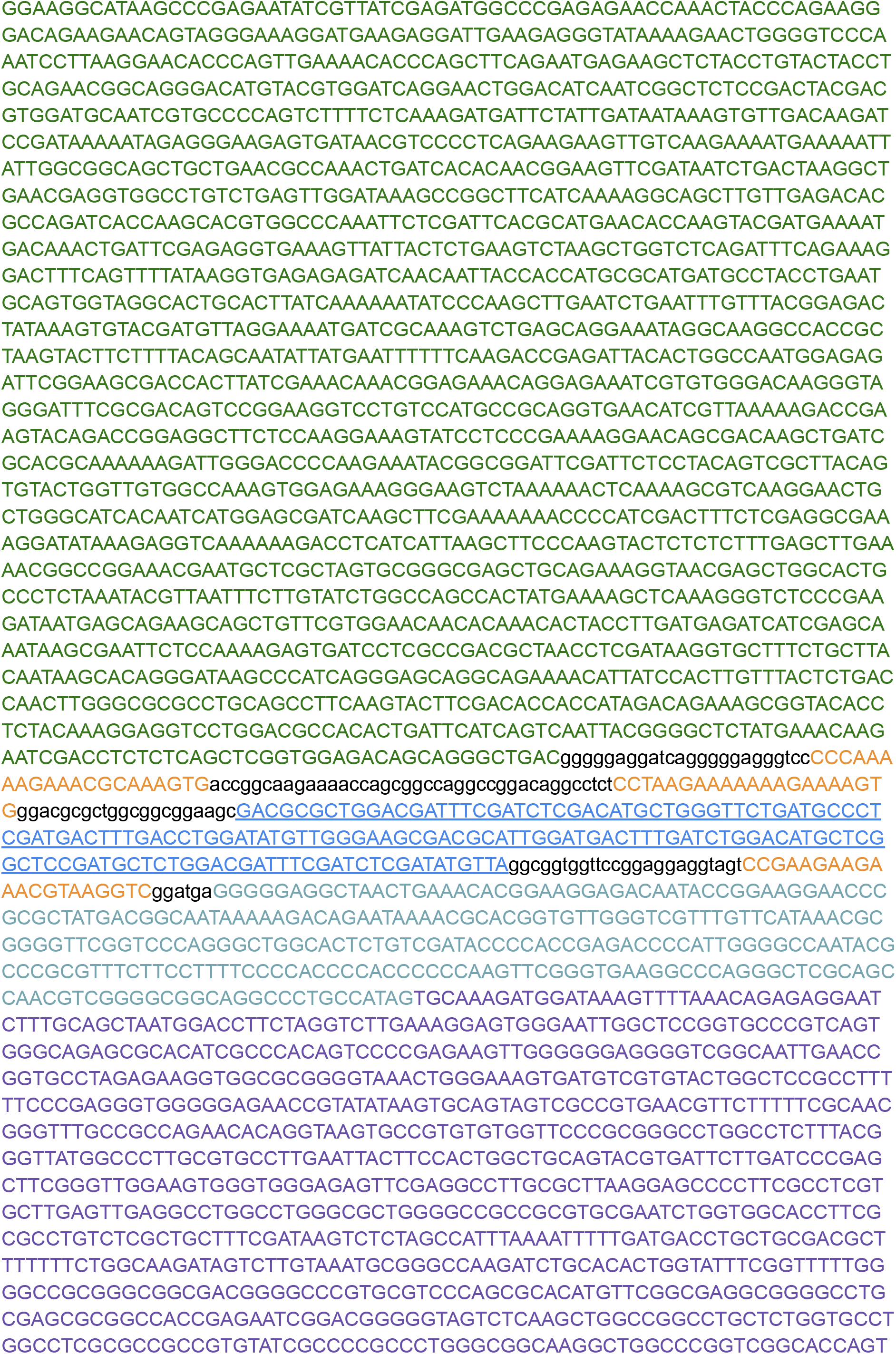
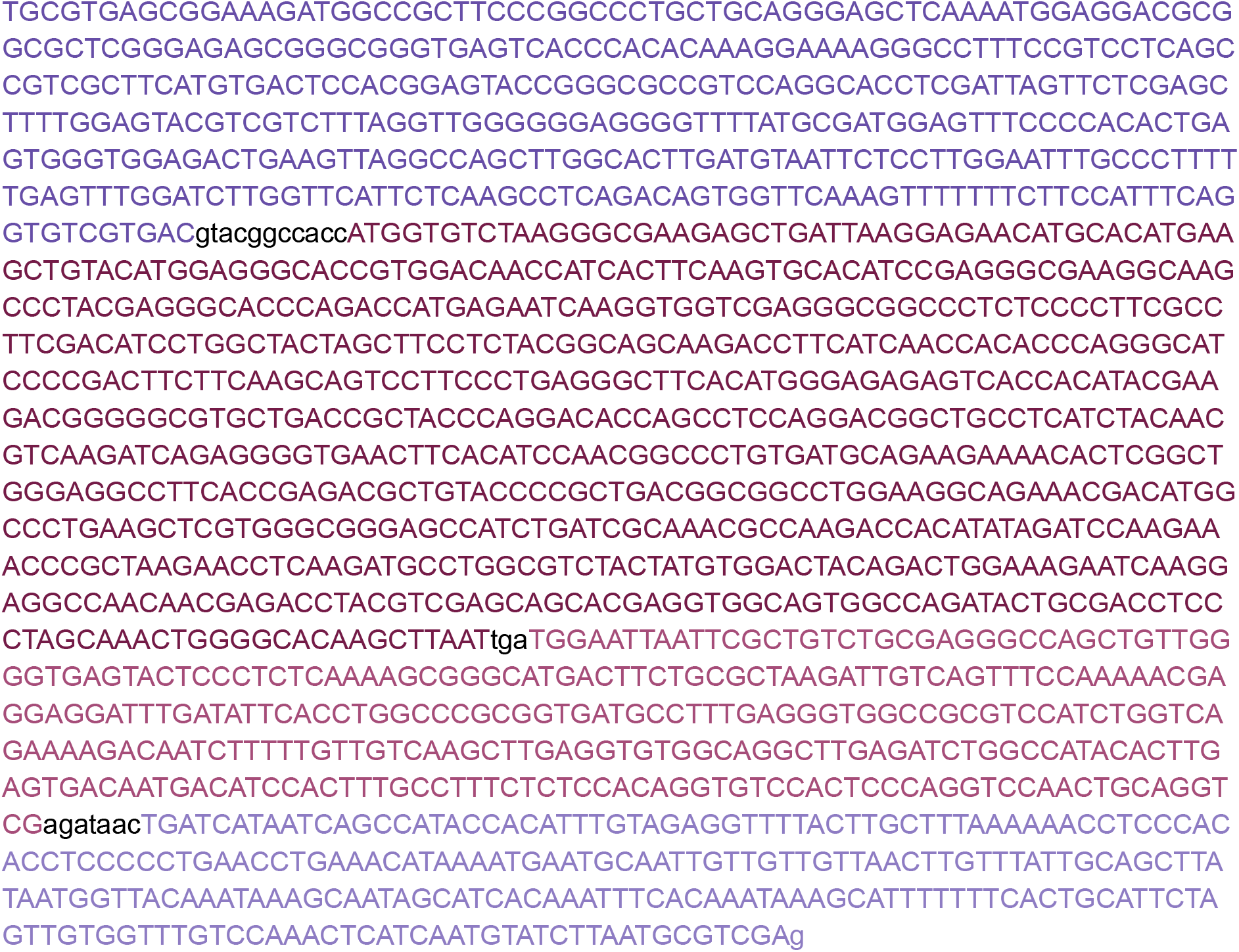

### Supp Info S2. SSP016_dCas9-p65HSF1

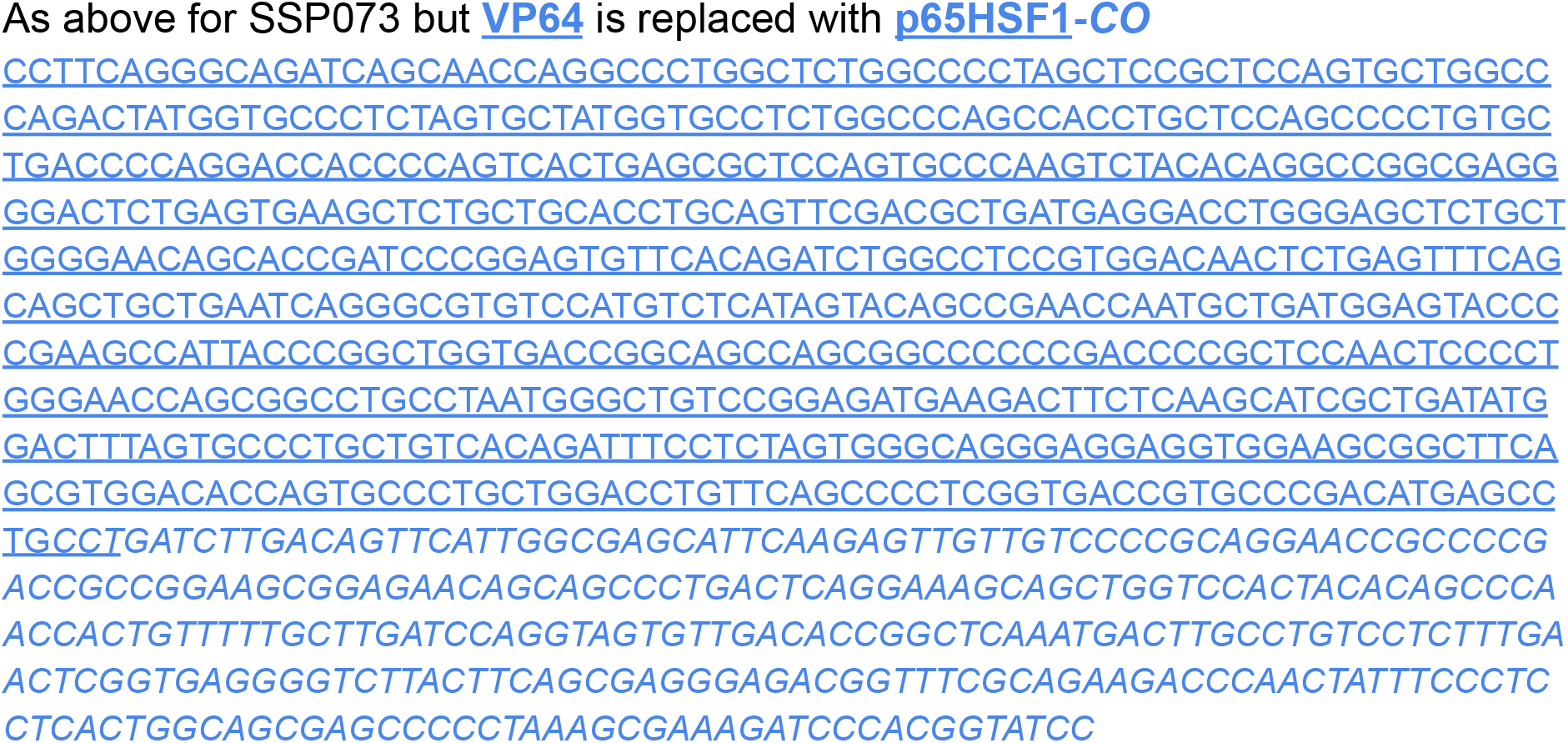

### Supp Info S3. SSP037_dCas9-VPR

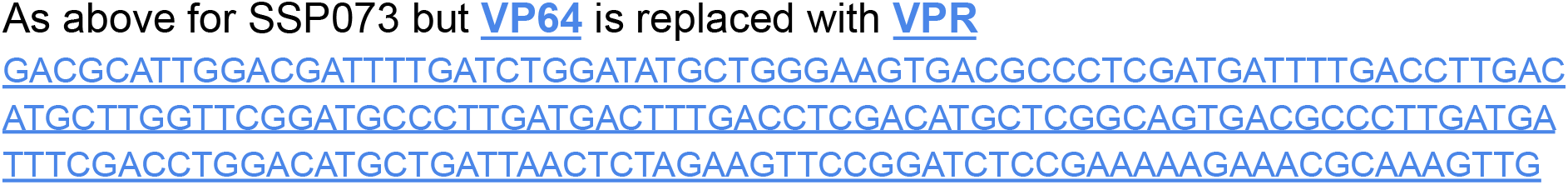
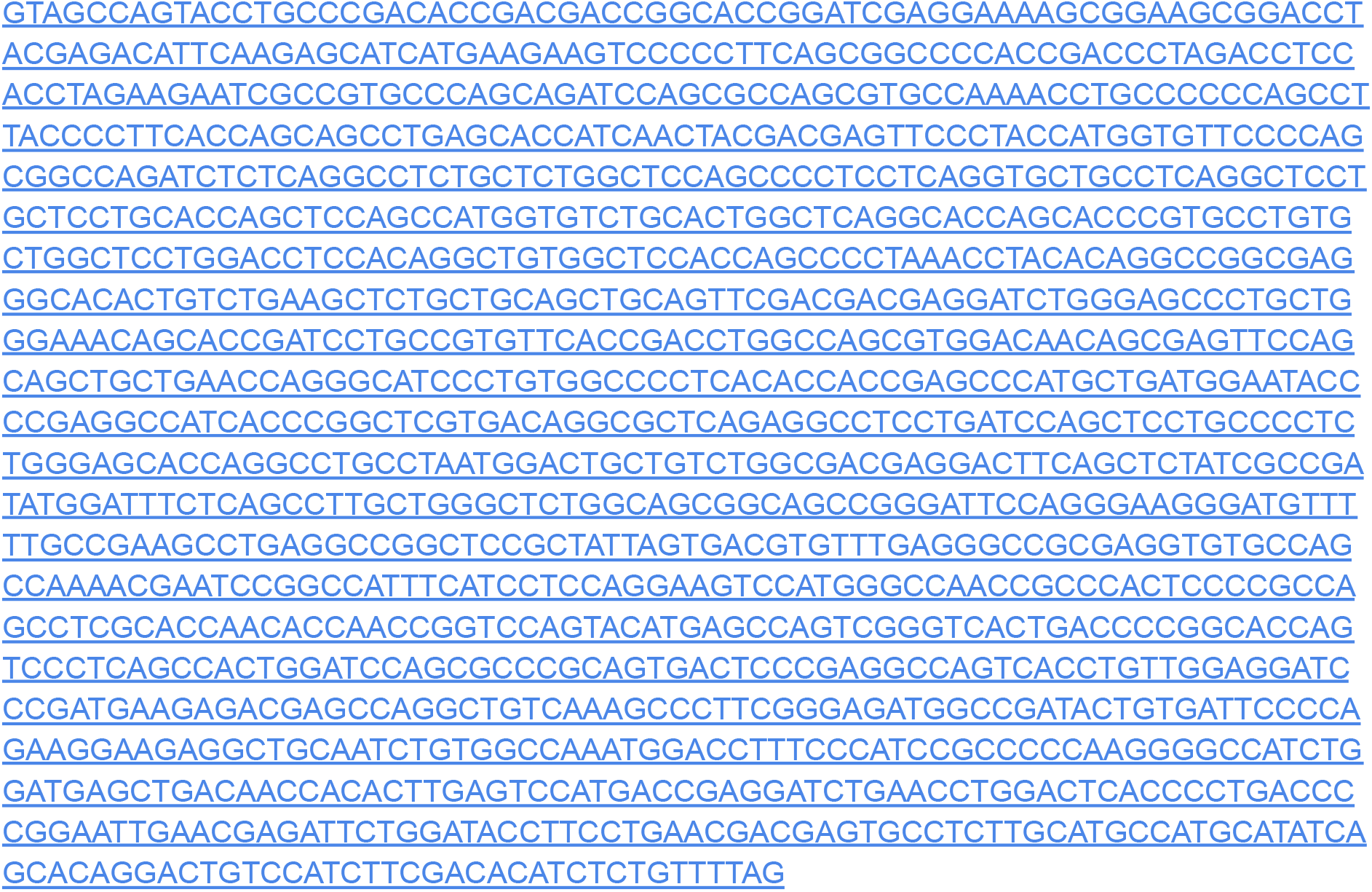

### Supp Info S4. SSP012_dCas9-SunTagx5-BFP

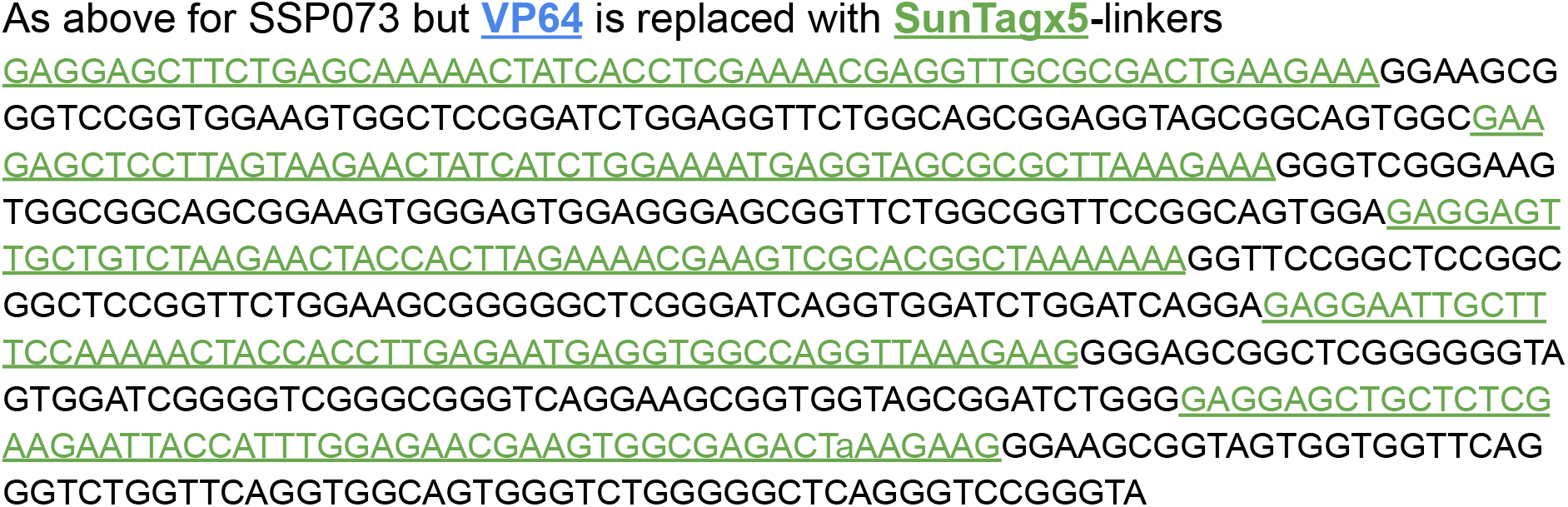

### Supp Info S5. SSP001_dCas9-SpyTagx1-BFP

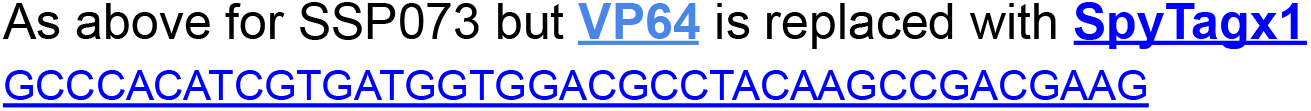

### Supp Info S6. SSP002_dCas9-SnoopTagx1-BFP

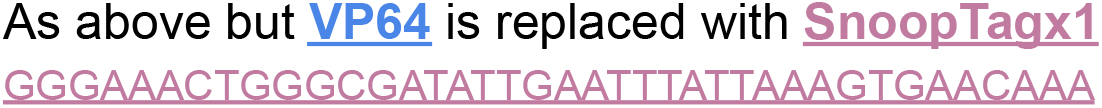

### Supp Info S7. SSP030_dCas9-SpyCatcherx1-BFP

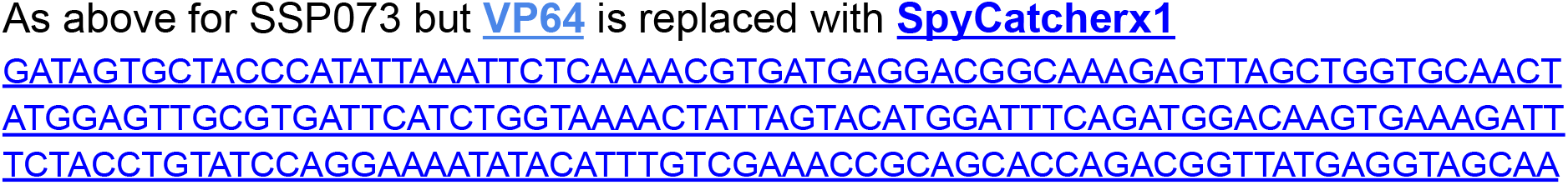
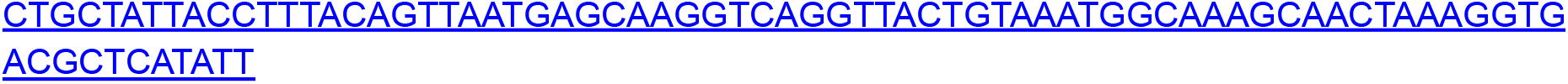

### Supp Info S8. SSP031_dCas9-SnoopCatcherx1-BFP

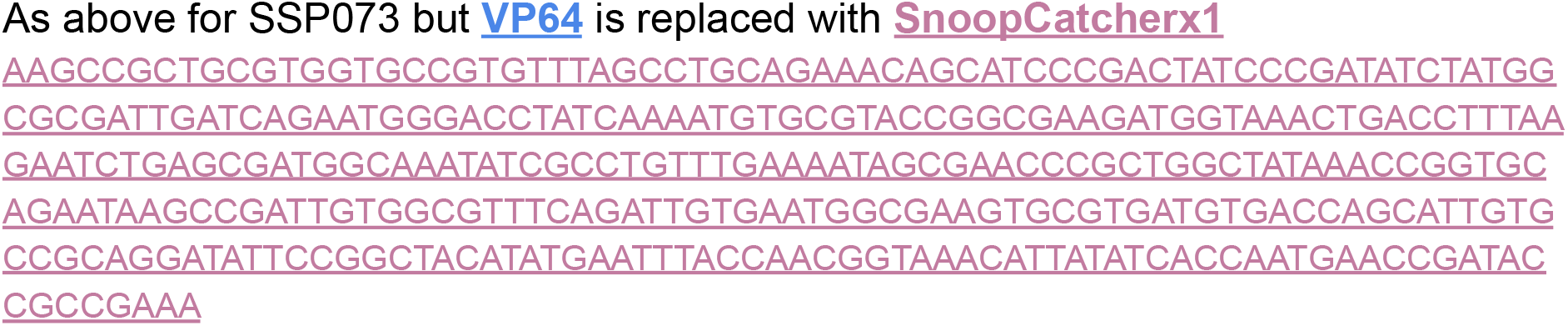

### Supp Info S9. SSP032_SpyTag-p65HSF1-CO

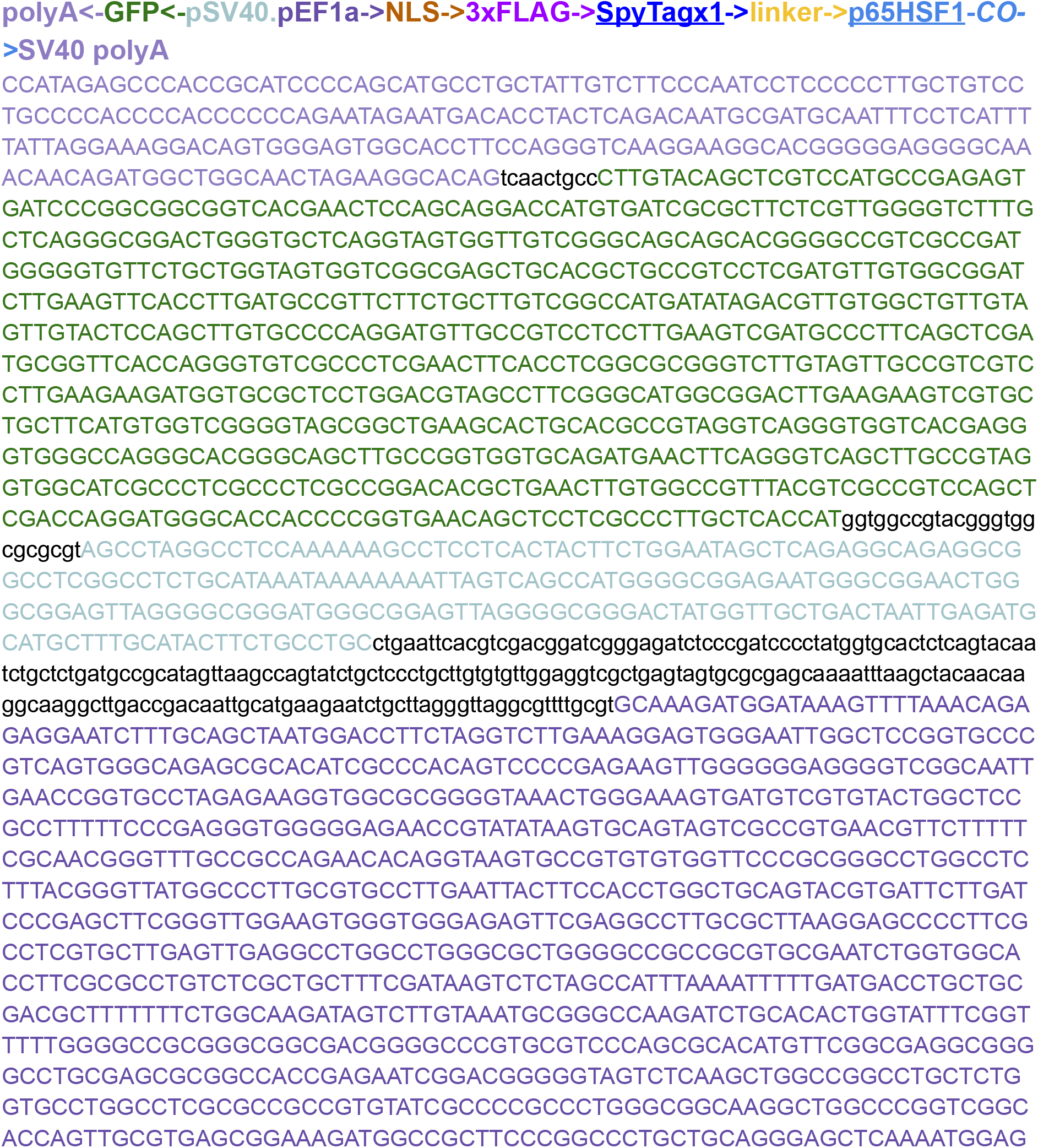
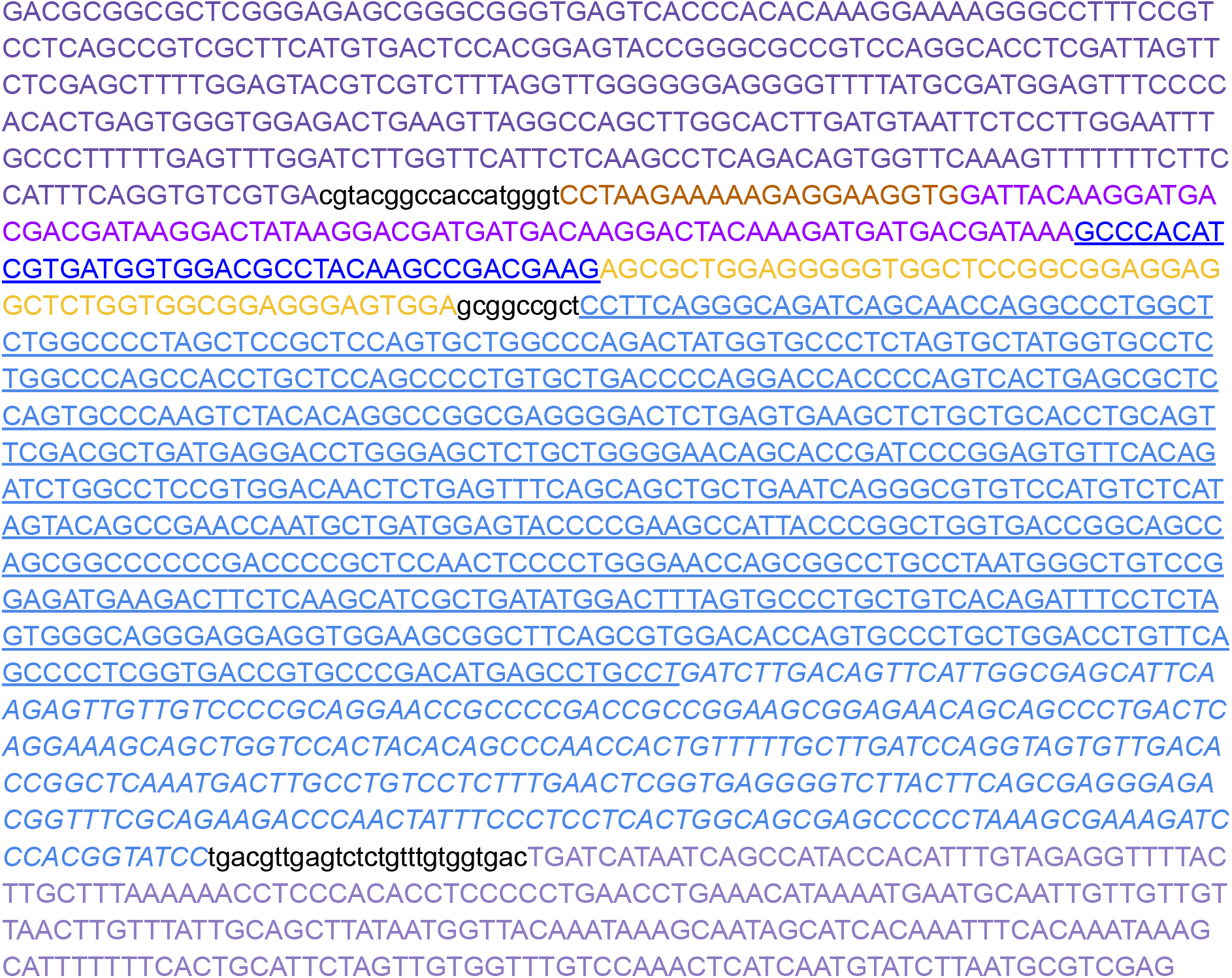

### Supp Info S10. SSP033_SnoopTag-p65HSF1-CO

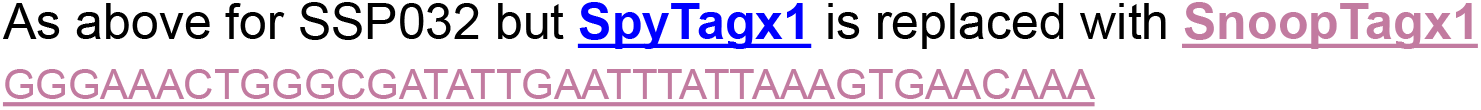

### Supp Info S11. SSP004_dCas9-Spy-Snoop-Sun-Avi-Tag-BFP

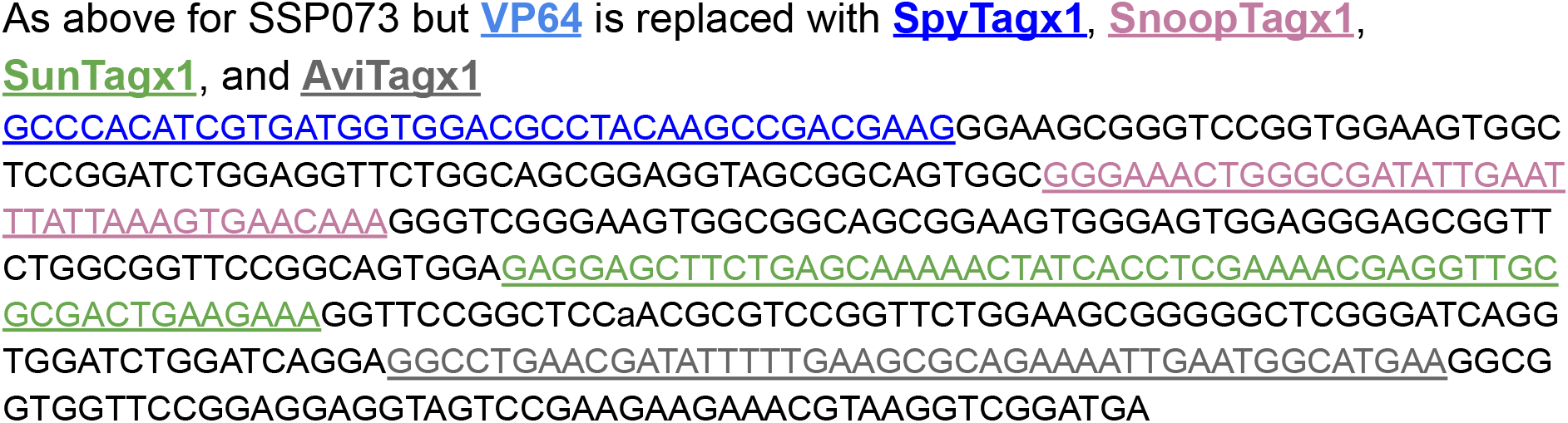

### Supp Info S12. SSP006_SpyCatcher-GFP-p65HSF1-CO

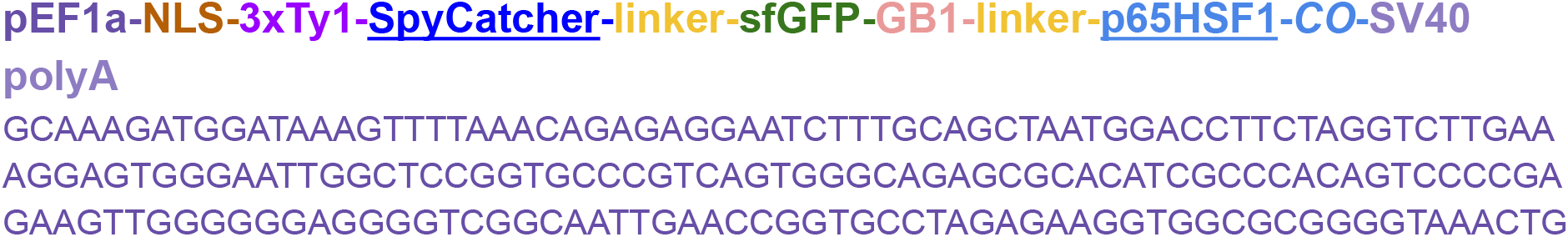
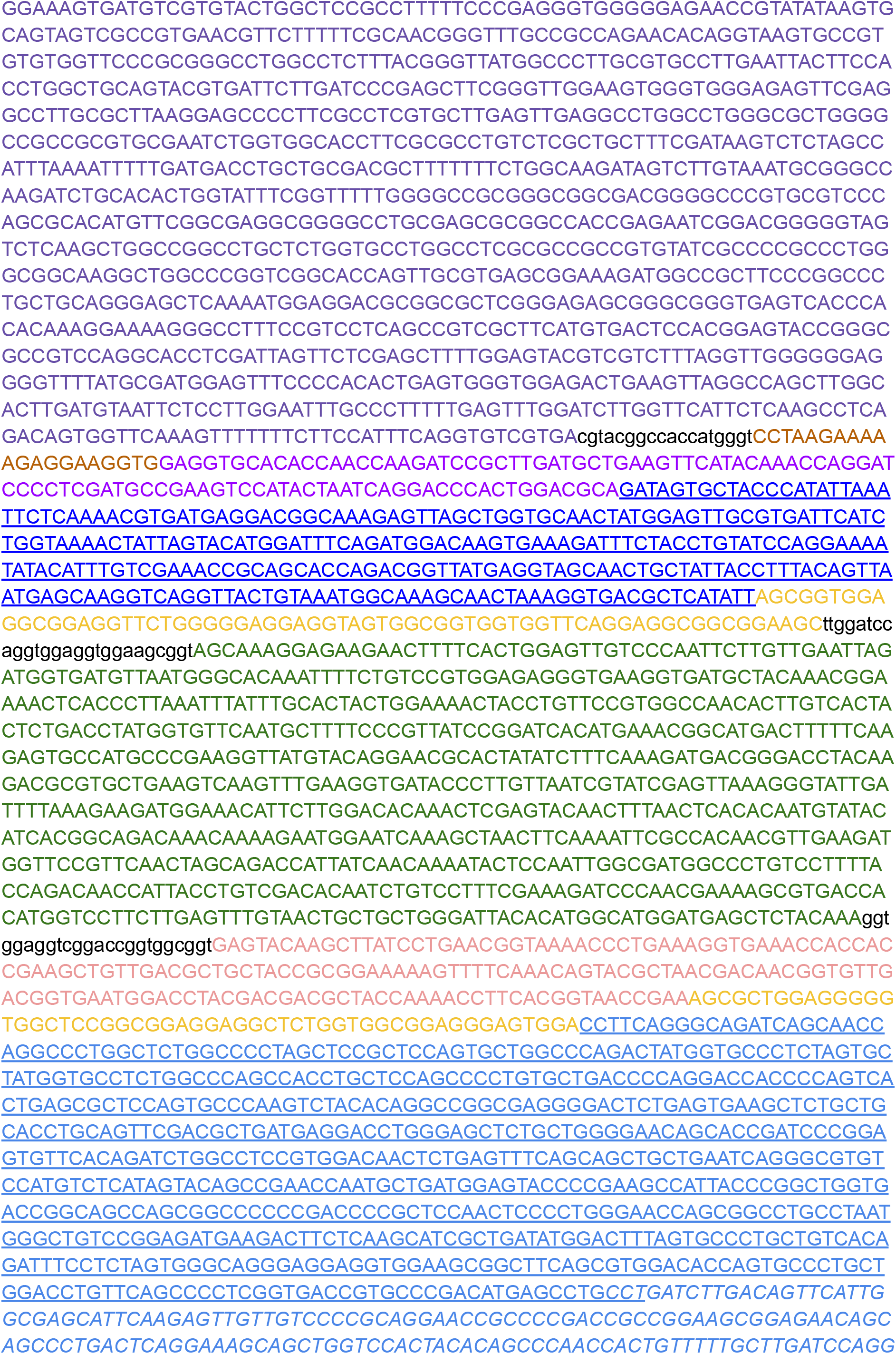
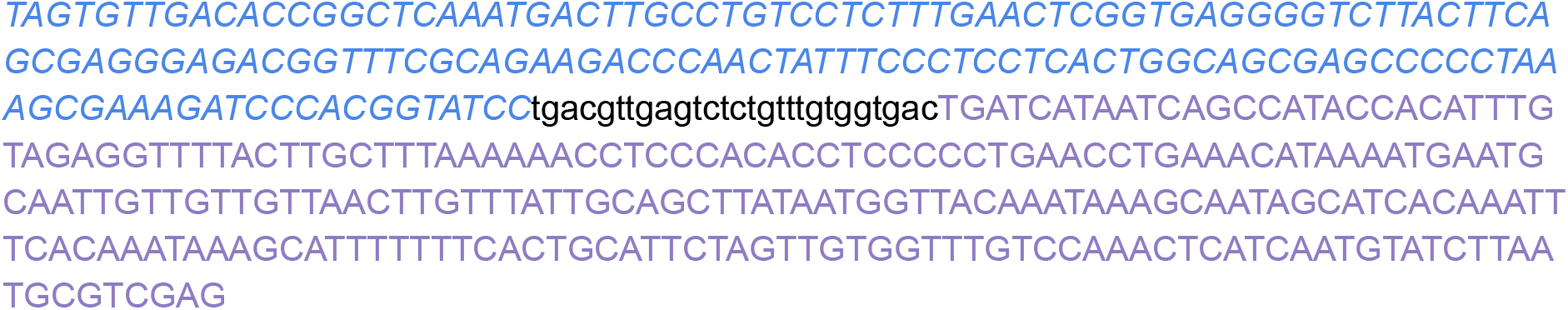

### Supp Info S13. SSP007_SpyCatcher-linker-p65HSF1-CO

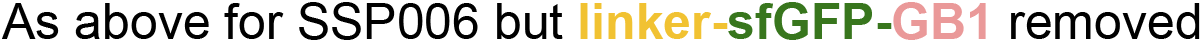

### Supp Info S14. SSP008_SnoopCatcher-GFP-p65HSF1-CO

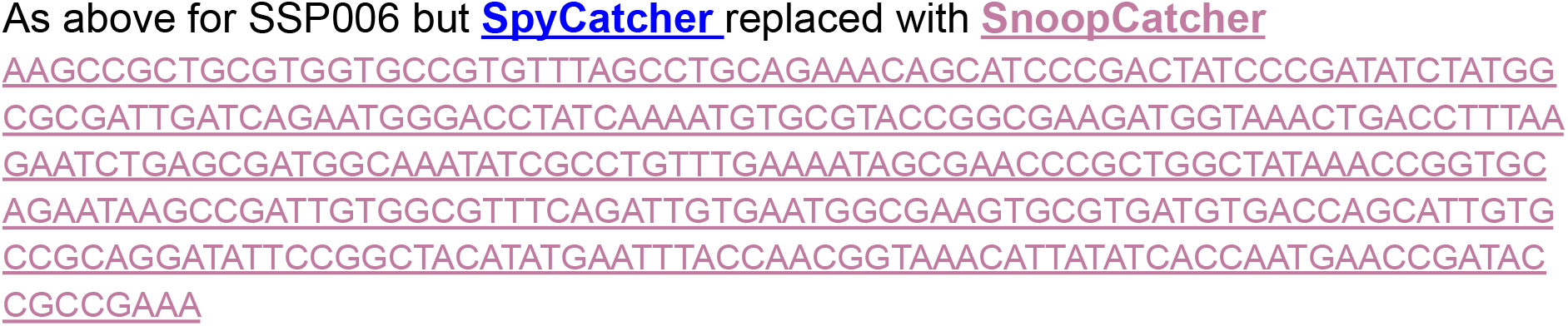

### Supp Info S15. SSP009_SnoopCatcher-linker-p65HSF1-CO

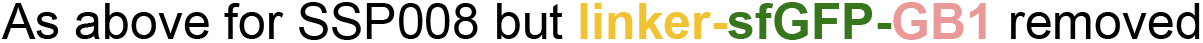

### Supp Info S16. SSP010_Traptavidin-GFP-p65HSF1-CO

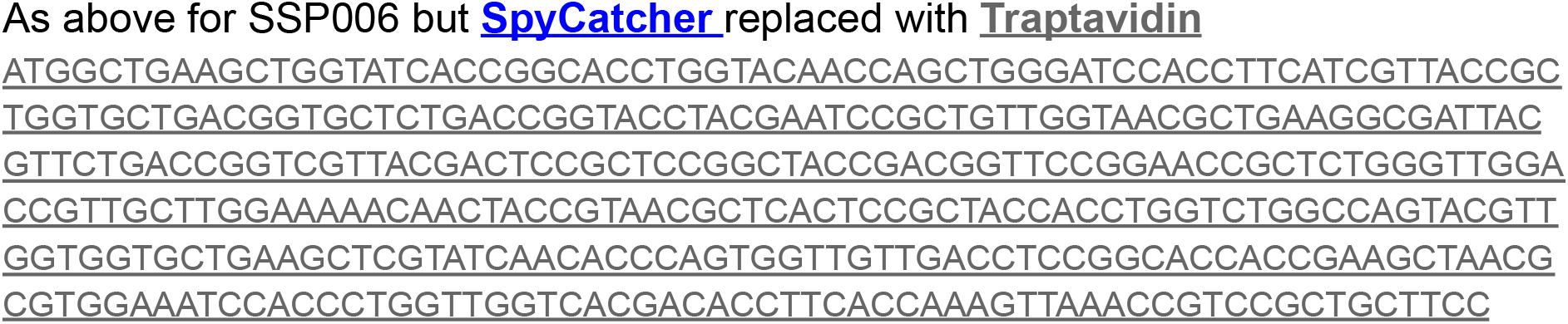

### Supp Info S17. SSP011_Traptavidin-linker-p65HSF1-CO

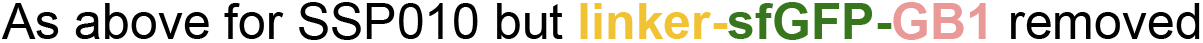

### Supp Info S18. SSP068_dCas9-SpyTagx5-BFP

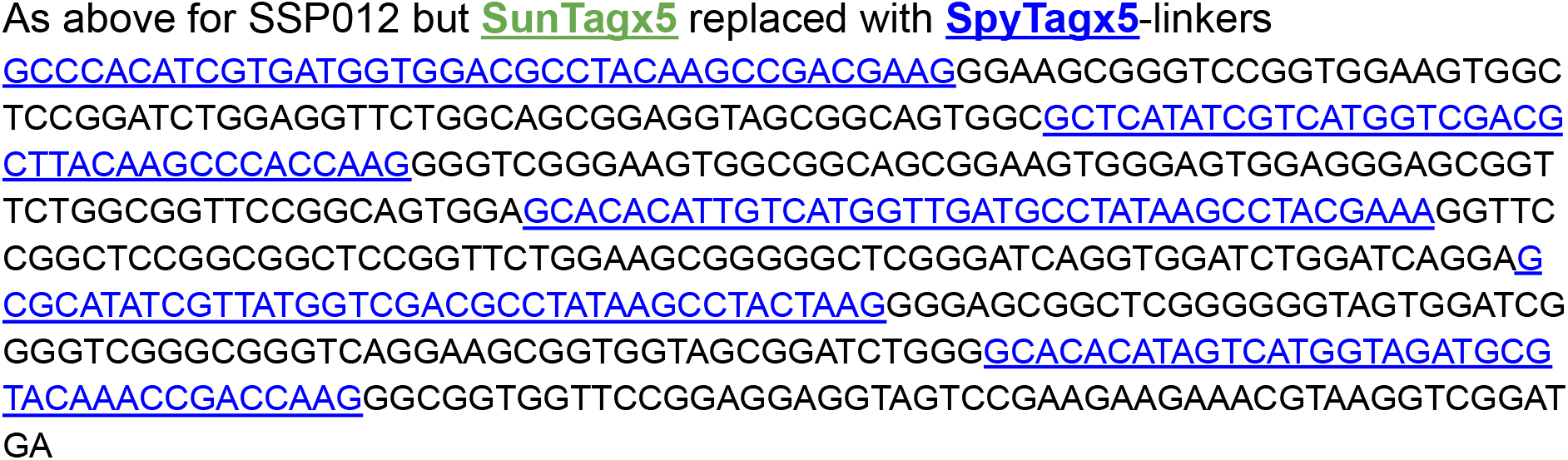

### Supp Info S19. SSP069_dCas9-SnoopTagx5-BFP

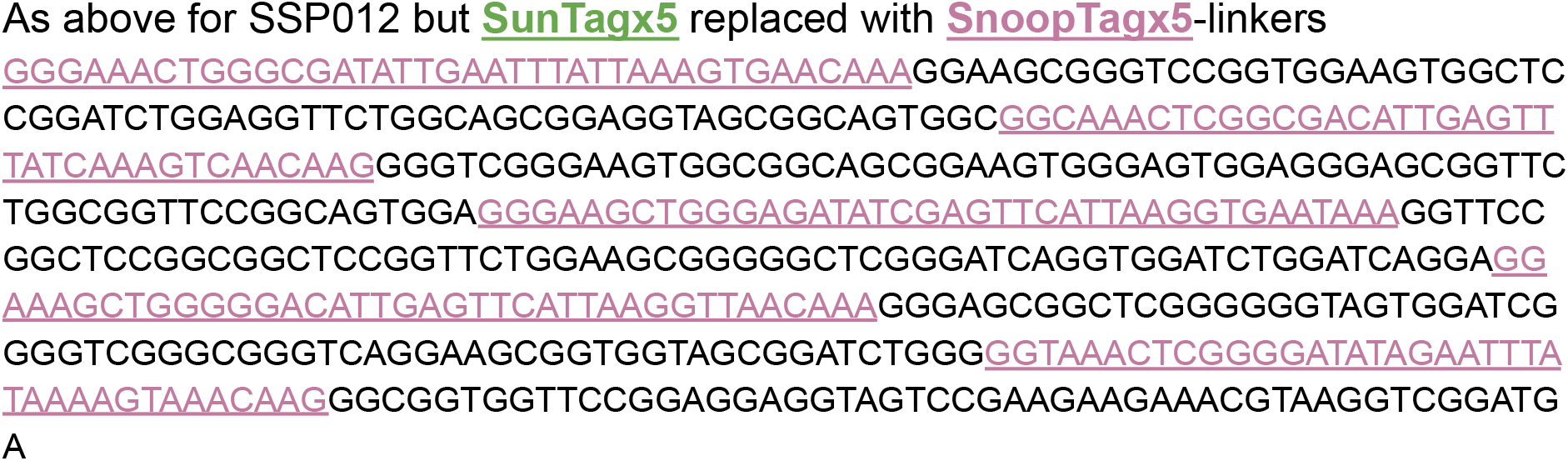

### Supp Info S20. SSP003_dCas9-AviTagx1-BFP

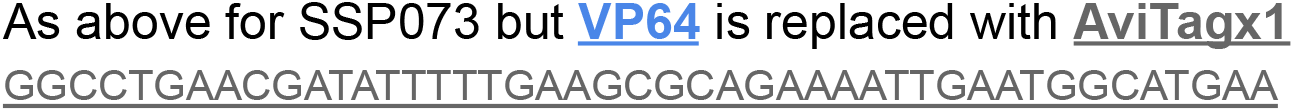

